# A partial reduction of Drp1 enhances mitophagy, autophagy, mitochondrial biogenesis, dendritic spines and synaptic activity in a transgenic Tau mouse model of Alzheimer disease

**DOI:** 10.1101/2021.10.07.463521

**Authors:** Ramesh Kandimalla, Maria Manczak, Jangampalli Adi Pradeepkiran, P. Hemachandra Reddy

**Author notes:** **Corresponding Author** P. Hemachandra Reddy, Ph.D., Department of Internal Medicine, Texas Tech University Health Sciences Center, 3601 4th Street, Lubbock TX 79430, United States, Lubbock, TX 79430, USA, Tel.: +1□806□743□3194.

## Abstract

The purpose of our study is to understand the impact of a partial dynamin-related protein 1 (Drp1) on cognitive behavior, mitophagy/autophagy, mitochondrial and synaptic activities in transgenic Tau mice in Alzheimer’s disease (AD). Our lab reported increased levels of Aβ and P-Tau, and abnormal interactions between Aβ and Drp1, P-Tau and Drp1 induced increased mitochondrial fragmentation and reduced fusion and synaptic activities in AD. These abnormal interactions, result in the proliferation of dysfunctional mitochondria in AD neurons. Recent research on mitochondria revealed that fission protein Drp1 is largely implicated in mitochondrial dynamics in AD. To determine the impact of reduced Drp1 in AD, we recently crossed transgenic Tau mice with Drp1 heterozygote knockout (Drp1+/-) mice and generated double mutant (Drp1+/- X Tau) mice. In the current study, we assessed cognitive behavior, mRNA and protein levels of mitophagy, autophagy, mitochondrial biogenesis, dynamics and synaptic genes, mitochondrial morphology & mitochondrial function, dendritic spines in Tau mice relative to double mutant mice. When compared to Tau mice, double mutant mice did better on Morris Maze (reduced latency to find hidden platform, increased swimming speed and time spent on quadrant) and rotarod (stayed a longer period of time) tests. Both mRNA and proteins levels autophagy, mitophagy, mitochondrial biogenesis and synaptic proteins were increased in double mutant mice compared to Tau mice. Dendritic spines were significantly increased; mitochondrial number is reduced and length is increased in double mutant mice. Based on these observations, we conclude that reduced Drp1 is beneficial in a symptomatic-transgenic Tau mice.

## Introduction

Alzheimer’s disease (AD) is a progressive, degenerative disorder manifested by dementia in aged individuals [1-4]. Over 50 million people worldwide are currently living with AD-related dementia, and this number is expected to increase to 152 million by 2050 (World Alzheimer Report 2019) [5]. AD-related dementia has huge economic consequences, as indicated by the total worldwide healthcare annual cost of dementia estimated in 2019 at $1 trillion. AD is associated with the loss of synapses, synaptic dysfunction, microRNA deregulation, abnormal redox signaling, mitochondrial structural and functional abnormalities, inflammatory responses, neuronal loss, accumulation of amyloid-beta (Aβ) and phosphorylated-tau (P-Tau) in disease progression [6-13].

Despite the progress that has been made in better understanding AD pathogenesis, researchers have still not identified early detectable markers, drugs, or agents that can prevent AD or slow its progression. Several decades of intense research have revealed that mitochondrial dysfunction and synaptic damage are early events in the disease process [14]. Mitochondrial function plays several key roles in synaptic maintenance [15,4], including assisting in ATP generation and Ca2+ regulation, both of which are essential for neuronal viability. Several studies found defective mitochondrial gene expression, increased free radical production, lipid peroxidation, oxidative DNA and protein damage, and decreased synaptic ATP production in postmortem AD brains, compared to healthy postmortem brains [15,16]. These changes primarily occur due to age-dependent increased production of -Tau and Aβ in neurons [4].

Further, using AD transgenic mice lines, studies have found increased production of free radicals and lipid peroxidation, and reduced levels of cytochrome c oxidase activity and synaptic ATP in AD-affected brain regions (cortex and hippocampus) [17-19], in the primary neurons of AD transgenic mice [15] and neurons expressing mutant APP and Aβ [20-24]. These findings strongly implicate mitochondrial dysfunction and oxidative stress in AD pathogenesis. Impaired mitochondrial dynamics, defective mitochondrial biogenesis, axonal transport of mitochondria, and synaptic damage are largely involved in mitochondrial dysfunction in AD [25, 4].

Maintaining mitochondrial integrity is an essential factor that contributes to the effective functioning of a cell. Therefore, any damaged/dysfunctional mitochondrial quality has a significant effect on neuronal health. Recent *in vitro* (APP hippocampal primary neurons) and *in vivo* studies (APP, APP/PS1, CRND8, and Tau mice) [26, 18, 19, 15, 27, 28, 22, 23, 20, 24, 29] revealed increased mitochondrial fragmentation, mitochondria dysfunction and defective mitophagy, strongly indicating that Aβ and P-Tau induced defective mitophagy in AD [30, 4]. Studies conducted in different AD models across species have revealed that abnormal mitochondria function, defective mitochondria dynamics, and compromised mitophagy lead to increased oxidative stress, synaptic dysfunction, neuronal loss, and cognitive decline, thereby contributing to enhanced AD pathologies [29, 26, 18, 19, 15]. Further, several studies have shown that mitophagy induction plays a protective role in ameliorating AD pathogenesis.

Recent studies on mitochondrial fission protein Drp1 knockout mice revealed that Drp1 homozygous knockouts are embryonic-lethal [31, 32]. However, heterozygote Drp1 knockout (Drp1+/-) mice are normal in terms of lifespan, fertility, and viability; and phenotypically, the Drp1+/- mice are not different from WT mice. To determine the effects of a partial reduction of Drp1 in the wild-type mice, we recently compared synaptic, dendritic, and mitochondrial proteins between Drp1+/- mice and wild-type mice and found Drp1+/- mice better than wild-type mice in all aspects [33].

Our lab reported increased levels of Aβ and P-Tau, and abnormal interactions between Aβ and Drp1, P-Tau and Drp1 and Aβ and P-Tau induced increased mitochondrial fragmentation and reduced mitochondrial fusion in AD [26, 33, 36]. These abnormal interactions result in the proliferation of dysfunctional mitochondria in AD neurons. Depleted Parkin and PINK1 levels occur due to increased accumulation of Aβ and P-Tau in the cytoplasm, reducing the effective number of effective autophagosomes targeted to dysfunctional mitochondrial and enhanced synaptic and mitochondrial fusion proteins relative to the wild-type mice, and the Drp1+/- mice showed significantly reduced free radicals and lipid peroxidation levels compared to the WT mice. These findings suggested that a partial reduction in Drp1 is beneficial with disease progression [26, 33, 36].

In studies to determine the protective effects of a partial reduction of Drp1 on Aβ and P-Tau, recently, we genetically crossed Drp1+/- mice with APP mice [34] and Drp1+/- mice with Tau mice [35], and then studied reduced Drp1 effects in 6-month-old APP and Tau mice. We found decreased mitochondrial fission and increased mitochondrial fusion, mitochondrial biogenesis, and synaptic proteins in the double mutant lines Drp1+/- X APP and Drp1+/- X Tau mice, suggesting that reduced free radicals and reduced excessive mitochondrial fragmentation may be protective against Aβ and P-Tau-induced toxicities in AD neurons. Therefore, it is important to study the protective effects of reduced Drp1 on the quality of mitochondria and mitophagy in 12-month-old mice (symptomatic condition).

In the current study, we assessed cognitive and motor coordination behaviors, dendritic spines, redox signaling/oxidative stress (free radicals and lipid peroxidation), mitochondrial function (ATP and cytochrome c oxidase activity), mRNA and protein levels of mitophagy, autophagy, mitochondrial biogenesis, dynamics and synaptic genes, mitochondrial morphology of reduced Drp1 in 12-month-old mutant Tau mice relative to Tau mice. The impact of reduced Drp1 on Tau pathologies and GTPase Drp1 activity are also assessed.

## Materials and methods

### Mice selection

To study the TauXDrp1effects, we used Drp1 heterozygote knockout mice (gifted by Hiromi Sesaki, Johns Hopkins University) and mutant Tau mice (P301L line) mice were purchased from Jackson Labs, New York, and used for our experiments.

### Dynamin-related protein 1 (Drp1) heterozygote knockout mice

Drp1+/− mice were generated using genetic a recombination strategy, replaced exons 3–5 of the GTPase domain, of the mouse endogenous Drp1 gene, with a neomycin-resistant gene (Wakabayashi et al 2009). Heterozygote Drp1 (+/−) mice are viable, fertile, normal in size, and do not show any phenotypic abnormalities. Heterozygous mice are maintained in a mixed background C57BL6/6-129/SvEv. We selected WT (+/+) or heterozygous (+/−) for Drp1 mice by genotype by using the DNA prepared from tail biopsy and PCR amplification Wakabayashi *et al*. (2009) [31].

### Mutant Tau mice

Tau P301L human mutation mice have purchased P301L mice from Taconic Farms (Cat # 1638-F, 1638-M) and used them for our investigations. These mice developed age-dependent hyperphosphorylated Tau and NFTs in the neocortex, hippocampus, and cortex. Cognitive impairments were found at 4.5 months of age in homozygous mice and 6 months of age in hemizygous mice. We did the genotyping for the human Tau P301L mutation by using the DNA prepared from tail biopsy and PCR amplification, as described in Lewis *et al*. (2001) [37].

### Double mutant (Drp1+/− X Tau) mice

We generated the double mutant (P301L, Drp1+/− X Tau) mice, by genetic crossing P301L mice with Drp1□+/− mice [38]. The resulting double mutant mice were studied for Tau pathology, mitochondrial and synaptic toxicities. We genotyped the Drp1+/− and P301L mutations, using DNA prepared from tail biopsy and for PCR amplification, as described in Wakabayashi *et al*. (2009) [31] and Lewis *et al*. (2000) [37]. All the mice were observed daily by a veterinary caretaker and further examined twice a week by laboratory staff, and if animals showed premature signs of neurological deterioration, they were euthanized before experimentation according to the procedure for euthanasia approved by the TTUHSC-IACUC and were not to be used in the proposed study.

### Quantitative real-time RT-PCR

Using the reagent TriZol (Invitrogen), total RNA was isolated from cortical tissues from Tau, Drp1+/−, Drp1+/− X Tau, and WT mice. Using primer Express Software (Applied Biosystems, Carlsbad, CA, USA), we designed the oligonucleotide primers for the housekeeping genes β-actin, GAPDH, mitochondrial structural genes, fission genes (Drp1, Fis1), fusion genes (MFN1, MFN2, Opa1), the mitochondrial matrix protein CypD, mitochondrial biogenesis genes PGC1α, PGC1β, Nrf1, Nrf2, TFAM and synaptic genes synaptophysin, PSD95, synapsins1-2, synaptobrevins1-2, neurogranin, GAP43, and synaptopodin, autophagy/mitophagy genes LC3B-I & II, LC3B, ATG5, Beclin1, PINK1, TERT, BCL2, and BNIP3L. The primer sequences and amplicon sizes are listed in **Table 1**. Using SYBR-Green chemistry-based quantitative real-time RT-PCR, we measured mRNA expression of the above-mentioned genes, as described by Kandimalla et al (2016) [38].

**Table 1.**
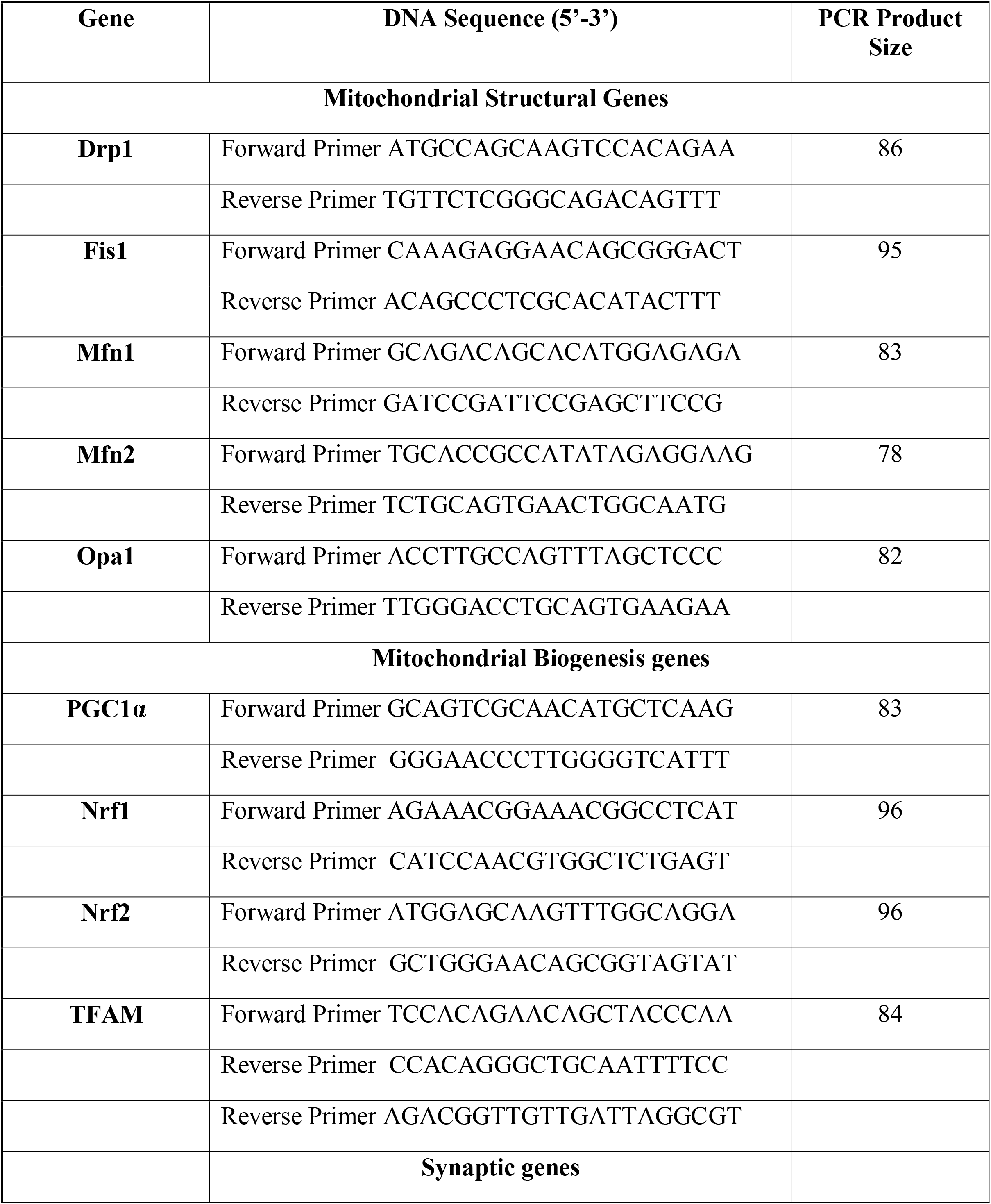
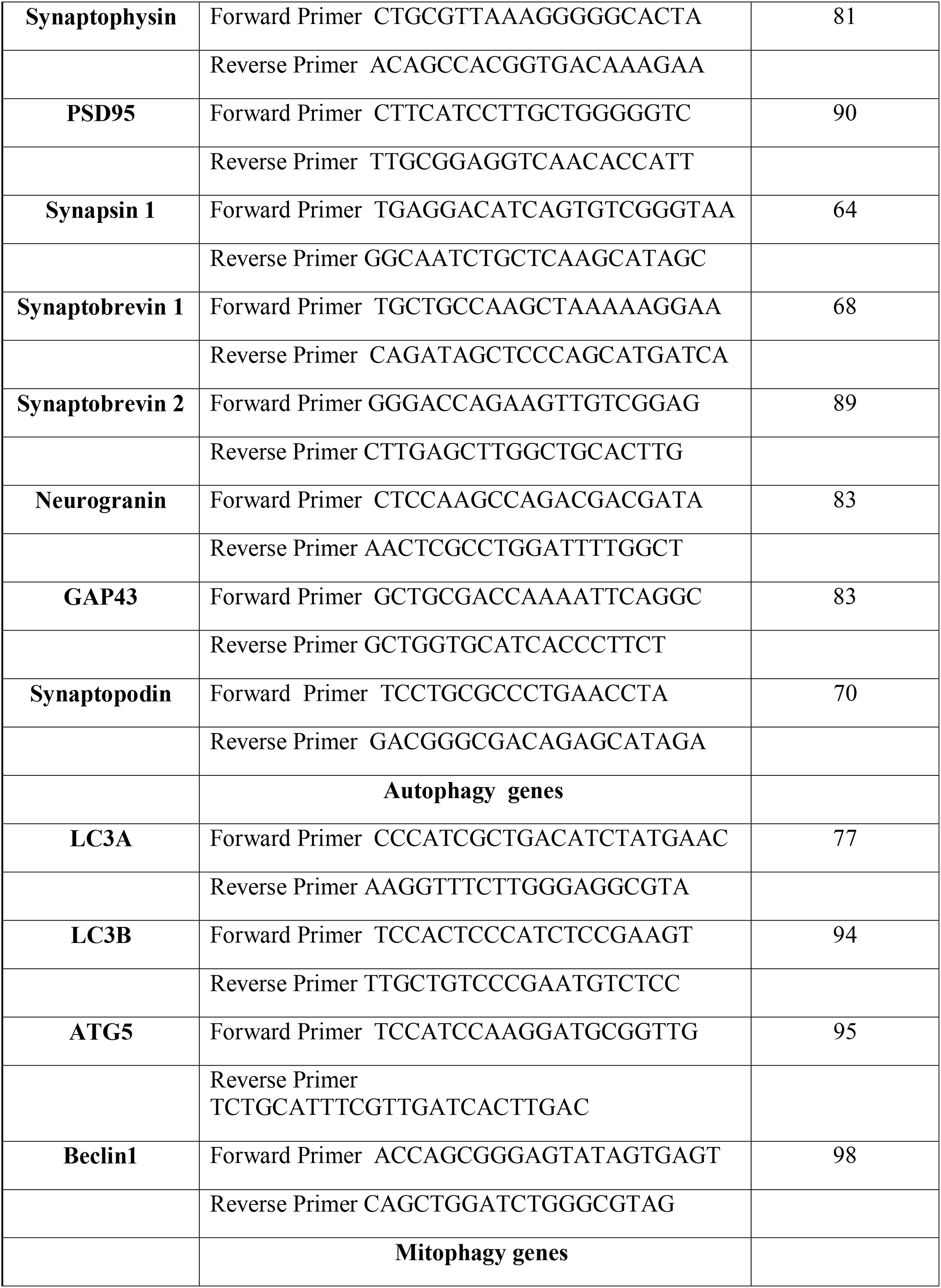
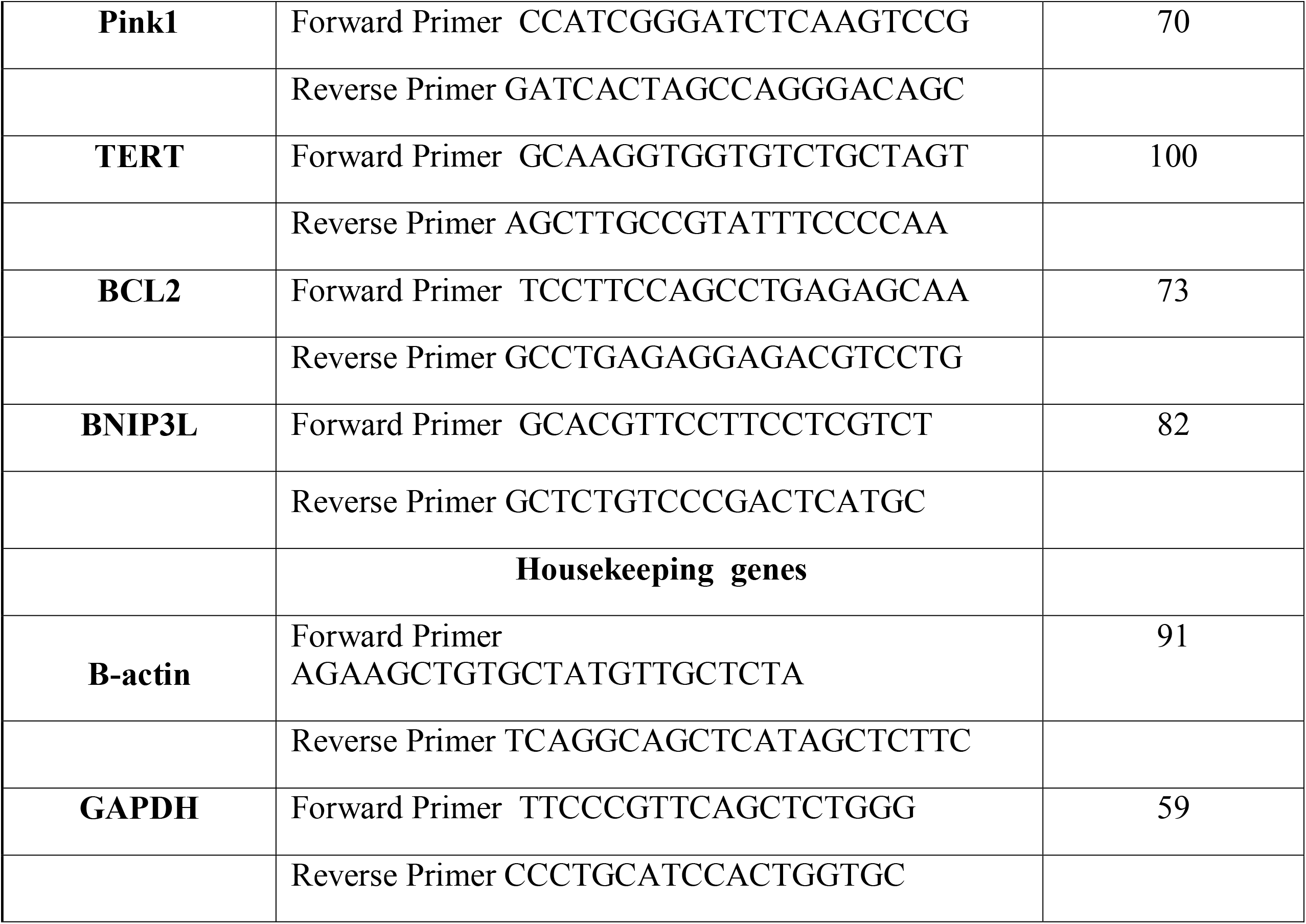
Summary of q RT-PCR oligonucleotide primers used in measuring mRNA expression in mitochondrial structural, biogenesis, synaptic genes and autophagy and mitophagy genes in WT, Drp1, Tau and Drp1-Tau in 12-month-old mice

### Immunoblotting analysis

To determine whether a partial deficiency of Drp1 alters the protein levels of mitochondrial dynamics, biogenesis, and synaptic genes that showed altered mRNA expression, we performed immunoblotting analyses of protein lysates from Tau, Drp1+/−, Drp1+/− X Tau and WT mice as described in Kandimalla et al 2016 [38]. Details of antibodies and their dilutions are given in **Table 2**.

**Table 2.** Summary of antibody dilutions and conditions used in the immunoblotting analysis of mitochondrial dynamics, biogenesisn autophagy and synaptic proteins.

| **Markers** | **Primary Antibody – Species and Dilution** | **Purchased from Company, City & State** | **Secondary Antibody, Dilution** | **Purchased from Company, City & State** |
| --- | --- | --- | --- | --- |
| DRP1 | Rabbit Polyclonal 1:300 | Novus Biological, Littleton, CO | Donkey Anti-rabbit HRP 1:10,000 | GE Healthcare Amersham, Piscataway, NJ |
| FIS1 | Rabbit Polyclonal 1:300 | Novus Biological, Littleton, CO | Donkey Anti-rabbit HRP 1:10,000 | GE Healthcare Amersham, Piscataway, NJ |
| Mfn1 | Rabbit Polyclonal 1:300 | Novus Biological, Littleton, CO | Donkey Anti-rabbit HRP 1:10,000 | GE Healthcare Amersham, Piscataway, NJ |
| Mfn2 | Rabbit Polyclonal 1:300 | Novus Biological, Littleton, CO | Donkey Anti-rabbit HRP 1:10,000 | GE Healthcare Amersham, Piscataway, NJ |
| OPA1 | Rabbit Polyclonal 1:300 | Novus Biological, Littleton, CO | Donkey Anti-rabbit HRP 1:10,000 | GE Healthcare Amersham, Piscataway, NJ |
| PGC1α | Rabbit Polyclonal 1:500 | Novus Biological, Littleton, CO | Donkey Anti-rabbit HRP 1:10,000 | GE Healthcare Amersham, Piscataway, NJ |
| Nrf1 | Rabbit Polyclonal 1:300 | Novus Biological, Littleton, CO | Donkey Anti-rabbit HRP 1:10,000 | GE Healthcare Amersham, Piscataway, NJ |
| Nrf2 | Rabbit Polyclonal 1:300 | Novus Biological, Littleton, CO | Donkey Anti-rabbit HRP 1:10,000 | GE Healthcare Amersham, Piscataway, NJ |
| TFAM | Rabbit Polyclonal 1:300 | Novus Biological, Littleton, CO | Donkey Anti-rabbit HRP 1:10,000 | GE Healthcare Amersham, Piscataway, NJ |
| SYN | Rabbit Monoclonal 1:400 | Novus Biological, Littleton, CO | Donkey Anti-rabbit HRP 1:10,000 | GE Healthcare Amersham, Piscataway, NJ |
| PSD95 | Rabbit Monoclonal<br>1:300 | Novus Biological,<br>Littleton, CO | Donkey Anti-rabbit HRP<br>1:10,000 | GE Healthcare<br>Amersham,<br>Piscataway, NJ |
| Phospho Tau<br>231 | Rabbit Polyclonal<br>1:200 | Thermo Fisher<br>Scientific, Waltham,<br>MA | Donkey anti-Mouse<br>1:10,000 | Thermo Fisher<br>Scientific,<br>Waltham, MA |
| Phospho Tau<br>181 | Rabbit Polyclonal<br>1:200 | Thermo Fisher<br>Scientific, Waltham,<br>MA | Donkey anti-Mouse<br>1:10,000 | Thermo Fisher<br>Scientific,<br>Waltham, MA |
| PINK1 | Rabbit Polyclonal<br>1: 1000<br>dilutions | Novus Biological,<br>Littleton, CO | Donkey Anti-rabbit HRP<br>1:10,000 | GE Healthcare<br>Amersham,<br>Piscataway, NJ |
| ATG5 | Rabbit Polyclonal<br>1: 1000<br>dilutions | Novus Biological,<br>Littleton, CO | Donkey Anti-rabbit HRP<br>1:10,000 | GE Healthcare<br>Amersham,<br>Piscataway, NJ |
| LCB-I | Rabbit Polyclonal<br>1: 1000<br>dilutions | Novus Biological,<br>Littleton, CO | Donkey Anti-rabbit HRP<br>1:10,000 | GE Healthcare<br>Amersham,<br>Piscataway, NJ |
| LCB-II | Rabbit Polyclonal<br>1: 1000<br>dilutions | Novus Biological,<br>Littleton, CO | Donkey Anti-rabbit HRP<br>1:10,000 | GE Healthcare<br>Amersham,<br>Piscataway, NJ |
| TERT | Rabbit Polyclonal<br>1: 1000<br>dilutions | Novus Biological,<br>Littleton, CO | Donkey Anti-rabbit HRP<br>1:10,000 | GE Healthcare<br>Amersham,<br>Piscataway, NJ |
| NDP52 | Mouse Polyclonal<br>1:500 | Novus Biological,<br>Littleton, CO | Donkey Anti-rabbit HRP<br>1:10,000 | GE Healthcare<br>Amersham,<br>Piscataway, NJ |
| Optineurin | Rabbit Polyclonal<br>1: 1000<br>dilutions | Novus Biological,<br>Littleton, CO | Donkey Anti-rabbit HRP<br>1:10,000 | GE Healthcare<br>Amersham,<br>Piscataway, NJ |
| P62 | Mouse Polyclonal<br>1:100 | Novus Biological,<br>Littleton, CO | Donkey Anti-rabbit HRP<br>1:10,000 | GE Healthcare<br>Amersham,<br>Piscataway, NJ |
| B-Actin | Mouse<br>Monoclonal<br>1:500 | Sigma-Aldrich,<br>St Luis, MO | Sheep Anti-mouse<br>HRP<br>1:10,000 | GE Healthcare<br>Amersham,<br>Piscataway, NJ |

### Mitochondrial functional assays

#### H_2_O_2_ production

Using an Amplex® Red H_2_O_2_ Assay Kit (Molecular Probes, Eugene, OR, USA), the production of H_2_O_2_ was measured using cortical tissues from Tau, Drp1+/−, Drp1+/− X Tau and WT mice as described in Kandimalla et al 2016 [38]. Hydrogen peroxide levels were compared between WT mice with Tau, Drp1+/−, double mutant (Drp1+/− X Tau) mice and data were also compared between Tau mice versus Drp1+/− X Tau mice.

#### Lipid peroxidation assay

Lipid peroxides are unstable indicators of oxidative stress in the brain. The final product of lipid peroxidation is HNE, which was measured in the cell lysates prepared from cortical tissues from Tau, Drp1+/−, Drp1+/− X Tau, and WT mice. We used HNE-His ELISA Kit (Cell BioLabs, Inc., San Diego, CA, USA) as described in Kandimalla et al 2016. Lipid peroxidation levels were compared between WT mice with Tau, Drp1+/−, double mutant (Drp1+/− X Tau) mice and data were also compared between Tau versus Drp1+/− X Tau mice.

#### Cytochrome oxidase activity

Cytochrome oxidase activity was measured in cortical tissues from all lines of mice. Enzyme activity was assayed spectrophotometrically using a Sigma Kit (Sigma–Aldrich) following the manufacturer’s instructions and as described in our lab publication by Kandimalla et al 2016 [38]. Cytochrome oxidase activity levels were compared between WT mice with Tau, Drp1+/−, double mutant (Drp1+/− X Tau) mice and data were also compared between Tau versus Drp1+/− X Tau mice.

#### ATP levels

ATP levels were measured in mitochondria isolated from cortical tissues of Tau, Drp1+/−, Drp1+/− X Tau, and WT mice and using an ATP determination kit (Molecular Probes), as described in Kandimalla et al 2016 [38]. ATP levels were compared between WT mice with Tau, Drp1+/−, double mutant (Drp1+/− X Tau) mice and data were also compared between Tau versus Drp1+/− X Tau mice.

#### GTPase Drp1 enzymatic activity

Using a colorimetric kit (Novus Biologicals, Littleton, CO, USA), GTPase enzymatic activity was measured in cortical tissues from Tau, Drp1+/−, Drp1+/− X Tau and WT mice following GTPase assay methods described in Kandimalla et al 2016 [38]. GTPase activity data were compared between WT mice with Tau, Drp1+/−, double mutant (Drp1+/− X Tau) mice and data were also compared between Tau mice versus Drp1+/− X Tau mice.

### Golgi-Cox staining and dendritic spine count

Golgi-Cox impregnation has been one of the most effective techniques for studying both the normal and abnormal morphology of neurons. The morphology of neuronal dendrites and dendritic spines have been discovered in the brains of mice by using Golgi-Cox staining and it was performed by using the FD Rapid GolgiStain Kit (FD Neuro Technologies, Columbia, MD, USA) as described in our lab publication by Kandimalla et al 2018 [19].

### Transmission electron microscopy

To determine the effects of phosphorylated tau on the mitochondrial number and size, we performed transmission electron microscopy in hippocampal and cortical sections of 12-month-old WT, Tau, Drp1+/- and double mutant (Drp1+/- X Tau) mice as described in our lab publications by Vijayan et al 2021. Electron microscopy was performed at 60 kV on a Philips Morgagni TEM equipped with a CCD, and images were collected at magnifications of□×1000–37 000. The numbers of mitochondria were counted and statistical significance was determined, using one-way ANOVA.

## Results

### Cognitive and motor coordination behavior

To determine the impact of reduced Drp1 on cognitive behavior in transgenic Tau mice, we performed multiple behavioral tests, including Rotarod and Morris Water Maze (latency to find the platform, time spent in the quadrant, swimming speed) and in 12-month-old non-transgenic wild type (WT) mice, transgenic Tau (P301L strain) mice, Drp1+/- mice, and double mutant (Drp1+/- X Tau) mice. We used 15 animals per group for all tests, WT mice (n= 15), 12-month-old Drp1+/- mice (n=15), 12-month-old Tau mice (n=15), and 12-month-double mutant Drp1+/- X Tau mice (n=15).

### Rotarod

In comparison to 12-month-old non-transgenic WT, Drp1+/-, and double mutant (Drp1 X Tau), and Tau mice did not stay too long on an accelerating rotarod test (P=0.001) compared to age-matched WT mice, and the latency to fall in all four trials were consistent in Tau mice (**Figure 1A**). However, in the presence of reduced Drp1 in Tau mice, animals stayed on for a longer period compared to Tau mice, and this is true in almost all cases. It appears that in the presence of reduced Drp1, the lost motor learning and coordination abilities in double mutant (Drp1+/- X Tau) mice were markedly reversed.

**Figure 1.**
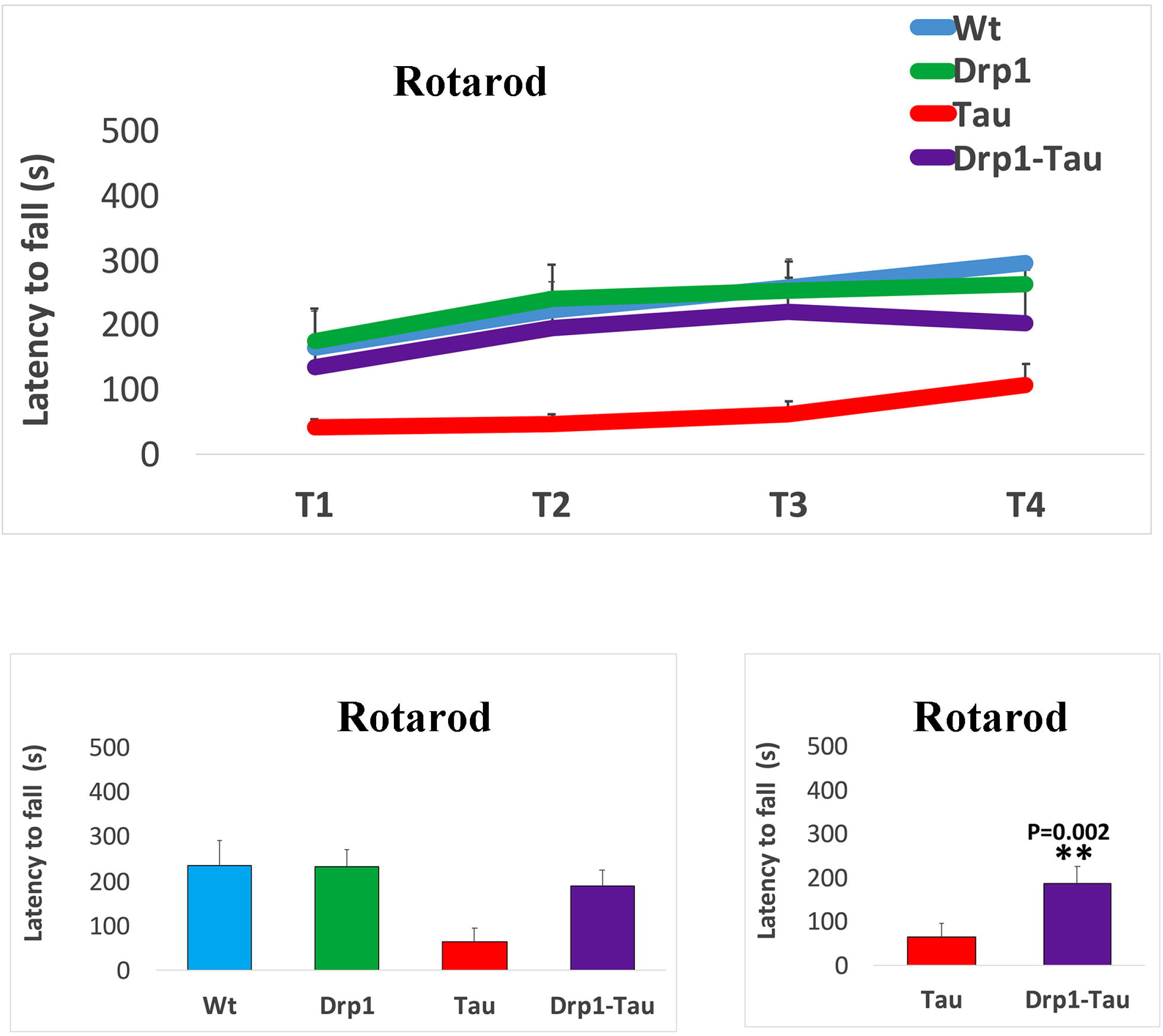
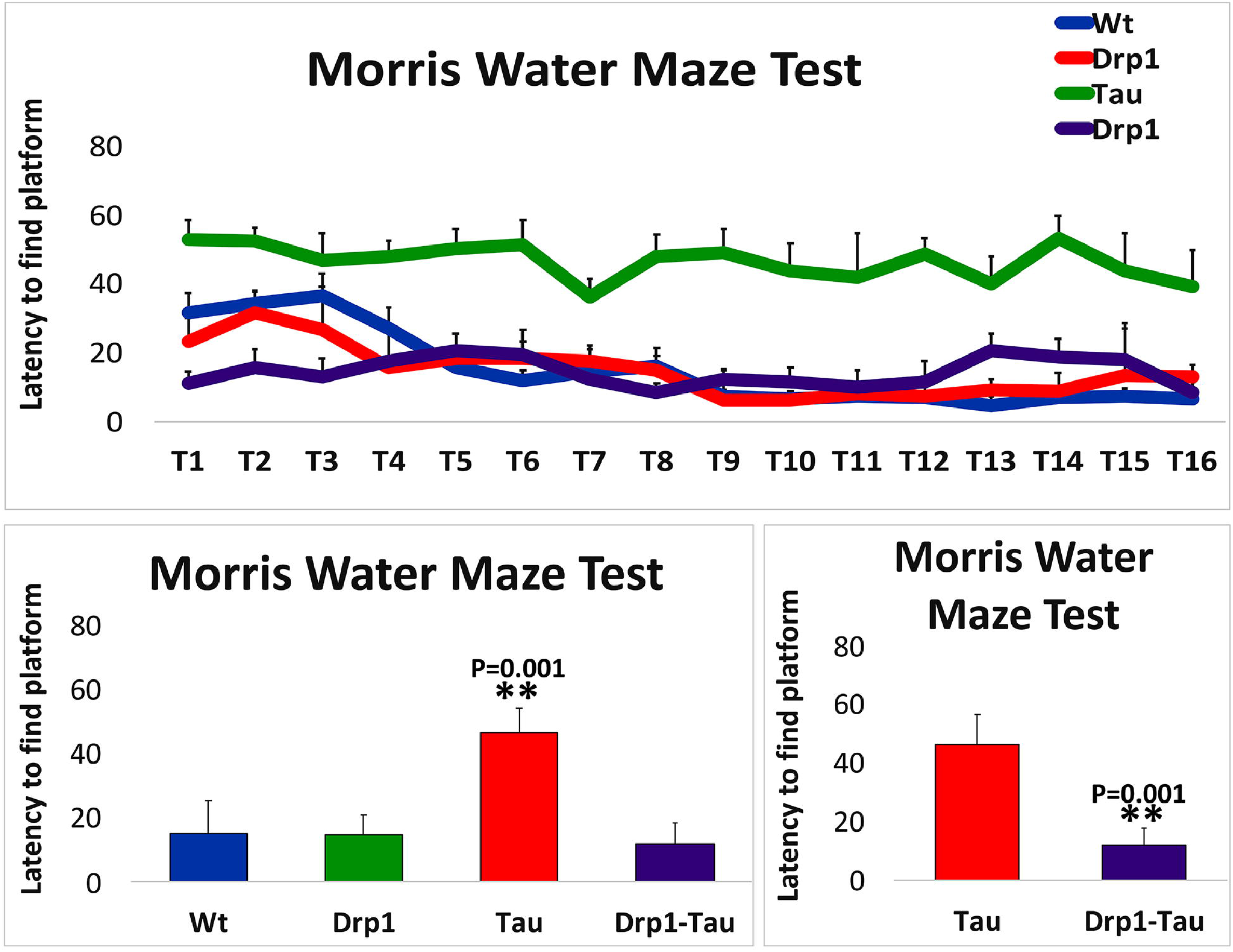
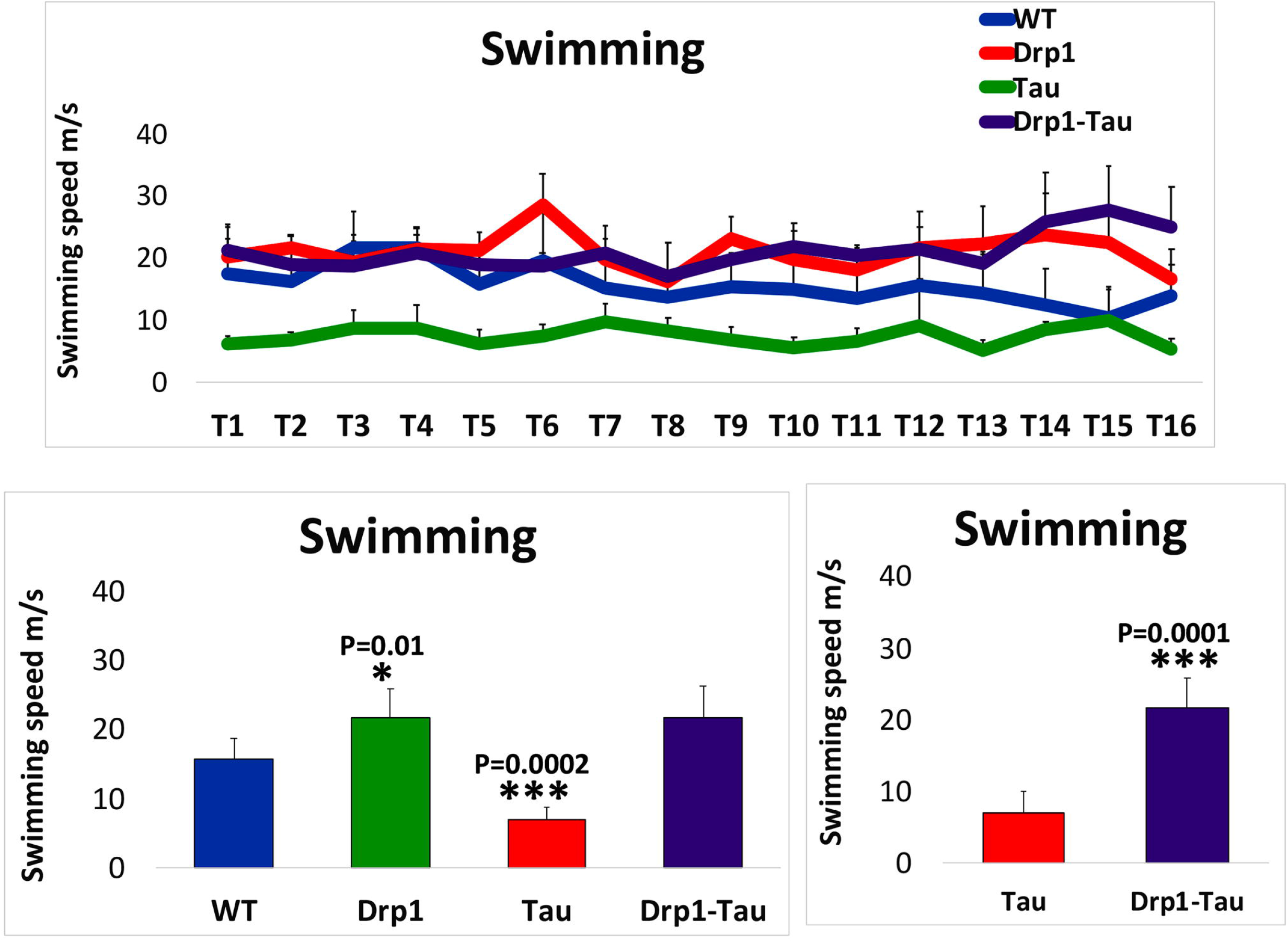
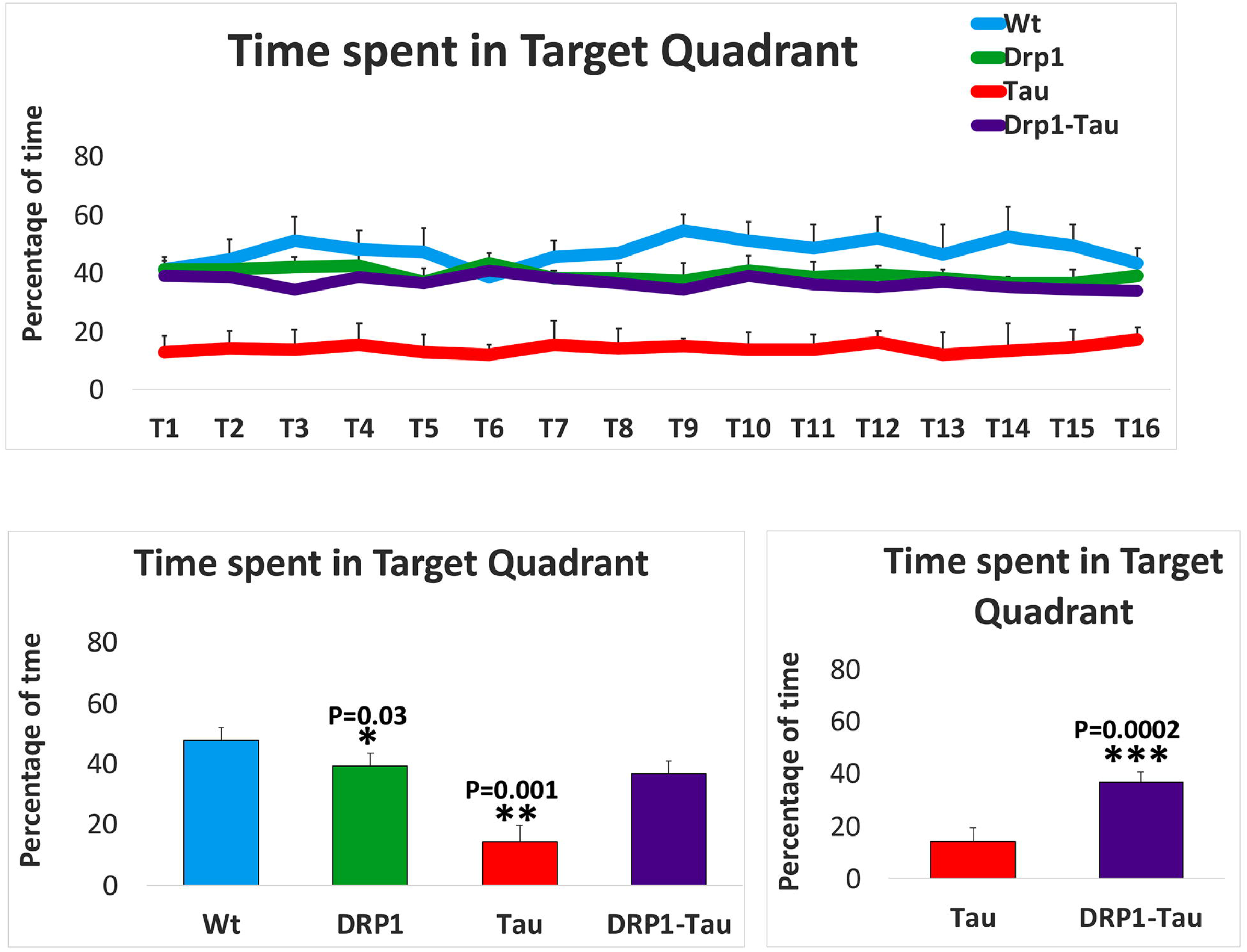
Cognitive behavior in 12-month-old non-transgenic wild-type (WT) mice, Tau (P301L), Drp1+/- mice, and double mutant (Drp1+/ X Tau) mice. **(A)** Rotarod test in 12-month-old Non-transgenic WT mice (N= 15), 12-month-old Drp1+/- mice (N= 15), 12-month-old Tau mice (N= 15), and 12-month-double mutant Drp1 X Tau mice (N= 15). Tau mice did not do well on rotarod compared to WT mice; on the other hand, double mutant Drp1 X Tau mice stayed for a longer period on a moving rotarod. **(B)** Latency to find platform test in 12-month-old non-transgenic WT mice, Drp1+/- mice, Tau mice, and double mutant Drp1 X Tau mice. As shown in Figure, Tau mice took a longer period to find a platform compared to WT mice. When compared to Tau mice, double mutant Drp1 X Tau mice reached the hidden platform earlier. **(C)** Swimming speed in 12-month-old non-transgenic WT mice, Drp1+/- mice, Tau mice, and double mutant Drp1-Tau mice. Swimming speed is significantly decreased for Tau mice, on the other hand, Drp1+/- mice had a higher swimming speed. When compared to Tau mice, double mutant Drp1 X Tau mice showed significantly higher swimming speed. **(D)** Time spent target quadrant in 12-month-old Non-transgenic WT mice, Drp1+/- mice, Tau mice, and double mutant Drp1-Tau mice. The average time spent in the target quadrant was significantly higher for double mutant mice compared to Tau mice.

### Morris Water Maze test

As shown in **Figure 1B**, Tau mice took a significantly longer time to identify the hidden platform (S) (P=0.001) than WT mice. However, double mutant (Drp1+/- X-Tau) mice, reached the hidden platform significantly earlier (P=0.001) than Tau mice. Overall, Drp1 deficiency in Tau mice resulted in a significant reduction in the time took to identify a hidden platform.

### Swimming speed

Swimming speed is significantly higher in 12-month-old Drp1+/- mice (P=0.01) compared to age-matched WT mice (**Figure 1C**). On the other hand, swimming speed is reduced in Tau mice (P=0.0002) relative to WT mice. In comparison to Tau mice, double mutant (Drp1+/- X Tau) mice (P=0001) showed significantly increased swimming speed.

### Time Spent in Target Quadrant

Tau mice spent reduced time (represented in percentage) in all 16 trials on <u>the</u> target quadrant compared with WT, Drp1+/-, and double mutant (Drp1+/- X Tau) mice (**Figure 1D**). The average time spent on the target quadrant was significantly decreased in Tau mice (P=0.001) compared with WT mice. The time spent in the target quadrant was significantly increased in double mutant (Drp1+/- X Tau) mice (P=0.0002) relative to Tau mice (**Figure 1D**).

### mRNA changes

To determine whether reduced Drp1 protect neurons against phosphorylated Tau-induced autophagy, mitophagy, mitochondrial and synaptic toxicities in AD, we measured mRNA levels of autophagy, mitophagy, mitochondrial biogenesis, dynamics, and synaptic genes in 12-month-old cortical/hippocampal tissues from Drp1+/−, Tau, Drp1+/− X Tau mice relative to age-matched WT mice. We also compared mRNA data between Tau and Drp1+/− X Tau mice, to understand the protective role of reduced Drp1 in mutant Tau-induced autophagy, mitophagy, mitochondrial and synaptic genes.

### Autophagy and mitophagy genes in Drp1+/− mice versus WT mice

mRNA expression levels were increased in ATG5, by 1.7-fold (P=0.05); in LC3B by 1.6-fold (P=0.05) and LC3B-I and Beclin 1 in 12-month-old Drp1+/- mice compared to age-matched WT mice (**Table 3**). Interestingly, mitophagy genes (PINK1 by 1.9-fold, P=0.05; TERT by 1.7-fold, P=0.05; BCL2 by 1.9-fold, P=0.05 and BNIP3L by 2.0-fold, P=0.05) were significantly increased in Drp1+/- mice relative to WT mice (**Table 3**).

**Table 3.**
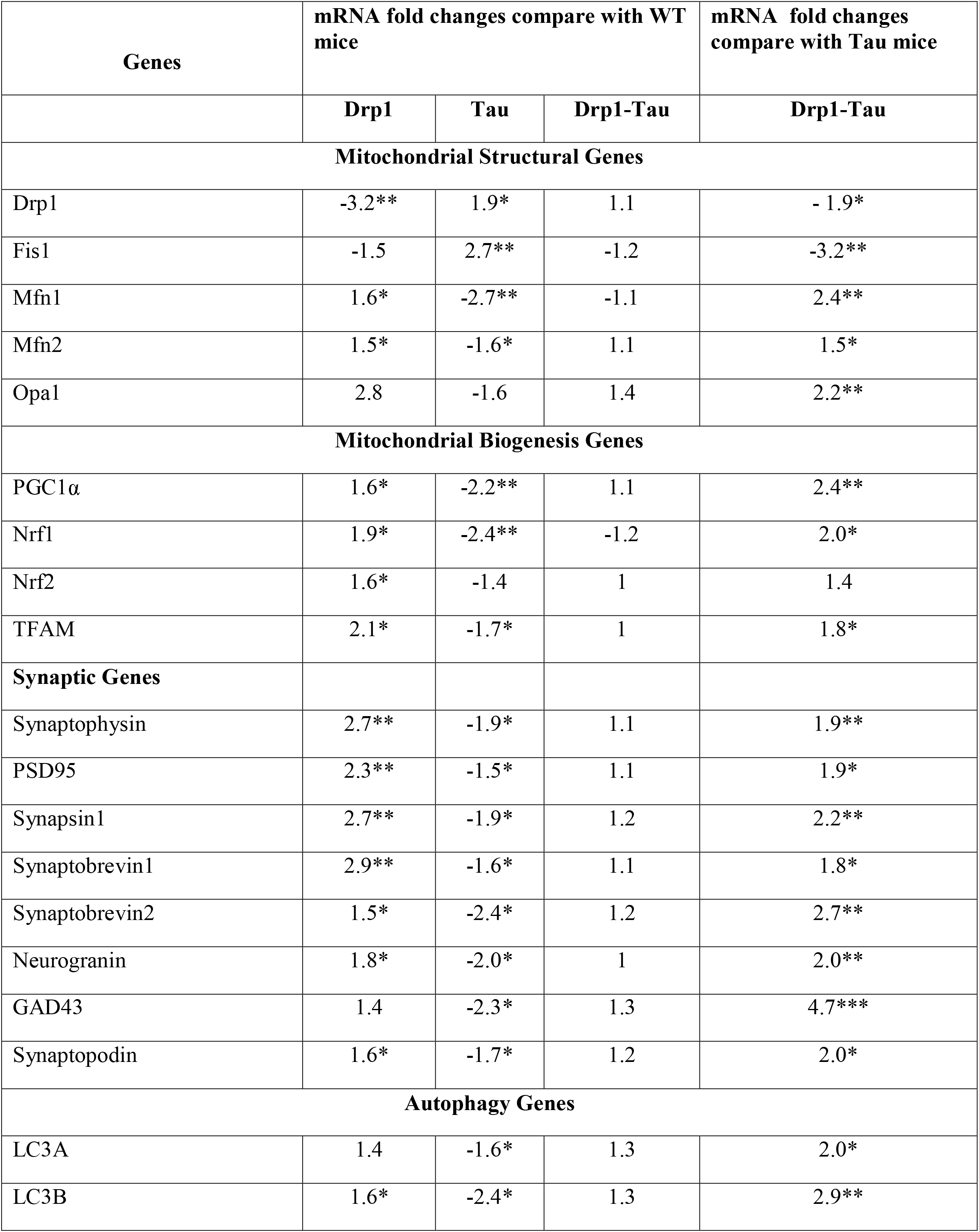
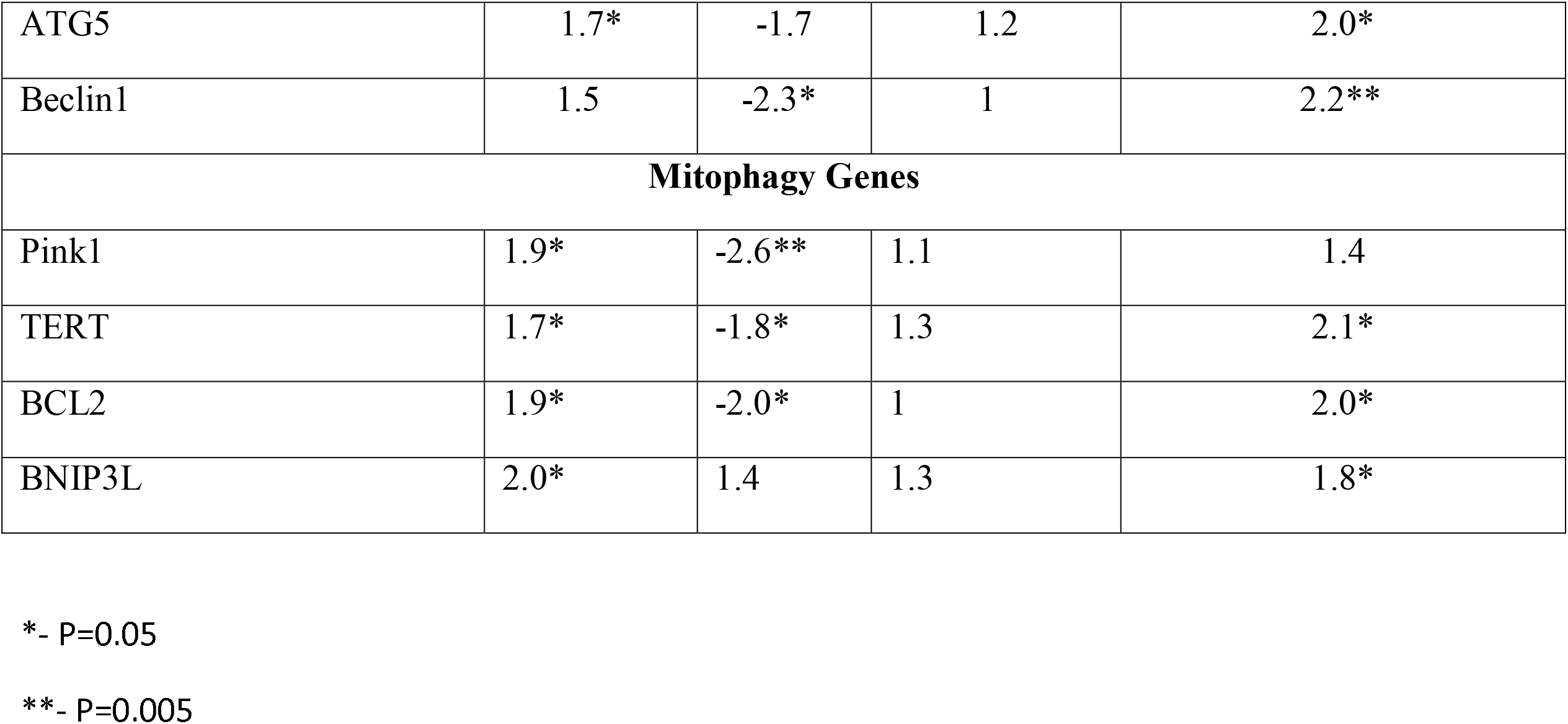
Fold changes of mRNA expression in mitochondrial dynamic, biogenesis, synaptic, autophagy and mitophagy related genes Drp1-Tau.

### Autophagy and mitophagy genes in Tau mice versus WT mice

In 12-month-old Tau mice compared to WT mice, mRNA expression levels were significantly reduced in autophagy genes ATG5, by 1.7-fold (*P*□=□0.05); in LC3BA by 1.6-fold, LC3B by 2.4-fold (P=0.05) and Beclin 1 by 2.3-fold and mitophagy genes PINK1 by 2.6-fold, P=0.005; TERT by 1.8-fold, P=0.05; BCL2 by 2.0-fold, P=0.05, and BNIP3L by 2.0-fold, P=0.05) (**Table 3**).

### Synaptic genes in Drp1+/- mice versus WT mice

mRNA expression levels were increased in synaptophysin, by 2.7-fold (*P*□=□0.005); in PSD95 by 2.3-fold (P=0.005); synapsin 1 by 2.7-fold, P=0.005; synaptobrevin 1 by 2.9-fold, P=0.005; synaptobrevin 1 by 1.5-fold, P=0.05; neurogranin by 1.8=fold, P=0.05; GAD43 by 1.4-fold and synaptopodin by 1.6-fold, P=0.05 in 12-month-old Drp1+/- mice compared to age-matched WT mice (**Table 3**).

### Synaptic genes in Tau mice versus WT mice

In 12-month-old Drp1+/- mice compared to age-matched WT mice, mRNA expression levels were reduced in synaptophysin, by 1.5-fold (P=0.05); in PSD95 by 1.5-fold (P=0.05); synapsin 1 by 1.9-fold, P=0.05; synaptobrevins 1 by 1.6-fold, P=0.05; synaptobrevins 2 by 2.4-fold, P=0.05; neurogranin by 2.0=fold, P=0.05; GAD43 by 2.3-fold and synaptopodin by 1.7-fold, P=0.05 in 12-month-old Drp1+/- mice compared to age-matched WT mice (**Table 3**).

### Mitochondrial biogenesis genes in Drp1+/- mice versus WT mice

Mitochondrial biogenesis genes (PGC1a by 1.6-fold, P=0.05; Nrf1 by 1.9-fold, P=0.05; Nrf2 by 1.6-fold and TFAM by 2.1fold, P=0.05) were significantly increased in 12-month-old Drp1+/- mice relative to age-matched WT mice (**Table 3**).

### Mitochondrial biogenesis genes in Tau mice versus WT mice

Mitochondrial biogenesis genes (PGC1a by 2.2-fold, P=0.005; Nrf1 by 2.4-fold, P=0.005; Nrf2 by 1.4-fold and TFAM by 1.7-fold, P=0.05) were significantly decreased in 12-month-old Drp1+/- mice relative to age-matched WT mice (**Table 3**).

### Mitochondrial dynamics in genes Drp1+/- mice versus WT mice

In 12-month-old Drp1+/- mice relative to age-matched WT mice, mitochondrial fission genes Drp1 by 3.2-fold, P=0.005; Fis1 by 1.5-fold were decreased. On the other hand, mitochondrial fusion genes Mfn1 by 1.6-fold, P=0.05; Mfn2 by 1.5-fold, P=0.05, and Opa1 by 2.8-fold were significantly increased in 12-month-old Drp1+/- mice relative to age-matched WT mice (**Table 3**).

### Mitochondrial dynamics in genes Tau mice versus WT mice

Mitochondrial fission genes Drp1 by 1.9-fold, P=0.05; Fis1 by 2.7-fold, P=0.005 were increased in 12-month-old Tau mice relative to age-matched WT mice (**Table 3**). On the other hand, mitochondrial fusion genes Mfn1 by 2.7-fold, P=0.005; Mfn2 by 1.6-fold, P=0.05, and Opa1 by 1.6-fold were significantly decreased in 12-month-old Tau mice relative to age-matched WT mice.

### Drp1+/- X Tau mice versus WT mice

As shown in **Table 3**, in 12-month-old Drp1+/- X Tau mice relative to age-matched WT mice, mRNA levels of autophagy, mitophagy, synaptic, mitochondrial biogenesis, and mitochondrial dynamics genes were not changed significantly.

### mRNA differences between Tau mice and Drp1+/− X Tau mice

We compared gene expression data between Drp1+/− X Tau and Tau mice, to understand whether a partial reduction of Drp1 protects against mutant Tau-induced autophagy, mitophagy, synaptic, mitochondrial biogenesis, and mitochondrial dynamics impairments.

#### Autophagy and mitophagy genes

In 12-month-old Drp1+/- X Tau mice relative Tau mice, mRNA expression levels of autophagy genes (ATG5, by 2.0-fold (P=0.05); in LC3A by 2.0-fold (P=0.05) and LC3B by 2.9-fold, P=0.005 and Beclin 1 by 2.2-fold, P=0.005) and mitophagy genes (TERT by 2.1-fold, P=0.05; BCL2 by 2.0-fold, P=0.05; BNIP3L by 1.8-fold, P=0.05, and PINK1 by 1.4-fold) were increased (**Table 3**).

#### Synaptic genes

mRNA expression levels were increased in synaptophysin, by 1.9-fold (P=0.005); in PSD95 by 1.9-fold (P=0.05); synapsin 1 by 2.2 -fold, P=0.005; synaptobrevin 1 by 1.8-fold, P=0.05; synaptobrevin 2 by 2.7-fold, P=0.005; neurogranin by 2.0=fold, P=0.005; GAP43 by 4.7-fold, P=0.0005 and synaptopodin by 2.0-fold, P=0.05 in 12-month-old Drp1+/- X Tau mice compared to age-matched Tau mice (**Table 3**).

#### Mitochondrial biogenesis genes

Mitochondrial biogenesis genes (PGC1α by 2.4-fold, P=0.005; Nrf1 by 2.0-fold, P=0.05; Nrf2 by 1.4-fold and TFAM by 1.8-fold, P=0.05) were significantly increased in 12-month-old Drp1+/-X Tau mice relative to age-matched Tau mice significantly increased in 12-month-old Drp1+/- X Tau mice relative to age-matched Tau mice (**Table 3**).

#### Mitochondrial dynamics genes

In 12-month-old Drp1+/- mice relative to age-matched WT mice, mitochondrial fission genes Drp1 by 1.9-fold, P=0.05; Fis1 by 3.2-fold, P=0.005 were decreased. On the other hand, mitochondrial fusion genes Mfn1 by 2.4-fold, P=0.005; Mfn2 by 1.5-fold, P=0.05, and Opa1 by 2.2-fold, P=0.005 were significantly increased in 12-month-old Drp1+/- X Tau mice relative to age-matched Tau mice (**Table 3**).

### Immunoblotting Analysis

#### Autophagy and Mitophagy proteins

To determine the protective effects of a partial reduction of Drp1 on autophagy and mitophagy proteins, we quantified autophagy and mitophagy proteins from cortical tissues of 12-month-old Drp1+/−, Tau, Drp1+/− X Tau, and WT mice.

##### Tau and WT mice

As shown in **Figure 2**, autophagy proteins ATG5, (P=0.02) LC3B-I (P=0.02), LC3B-II (P=0.02) and P62, LC3-binding adaptor protein (P=0.004), and TERT (autophagy inducer) (P=0.001) were significantly decreased in 12-month-old Tau mice relative to age-matched WT mice. Mitophagy proteins, PINK1 (P=0.004), Optineurin (P=0.004), and NDP52 (0.001) (mitophagy receptor proteins) were also significantly reduced in Tau mice relative control WT mice.

**Figure 2.**
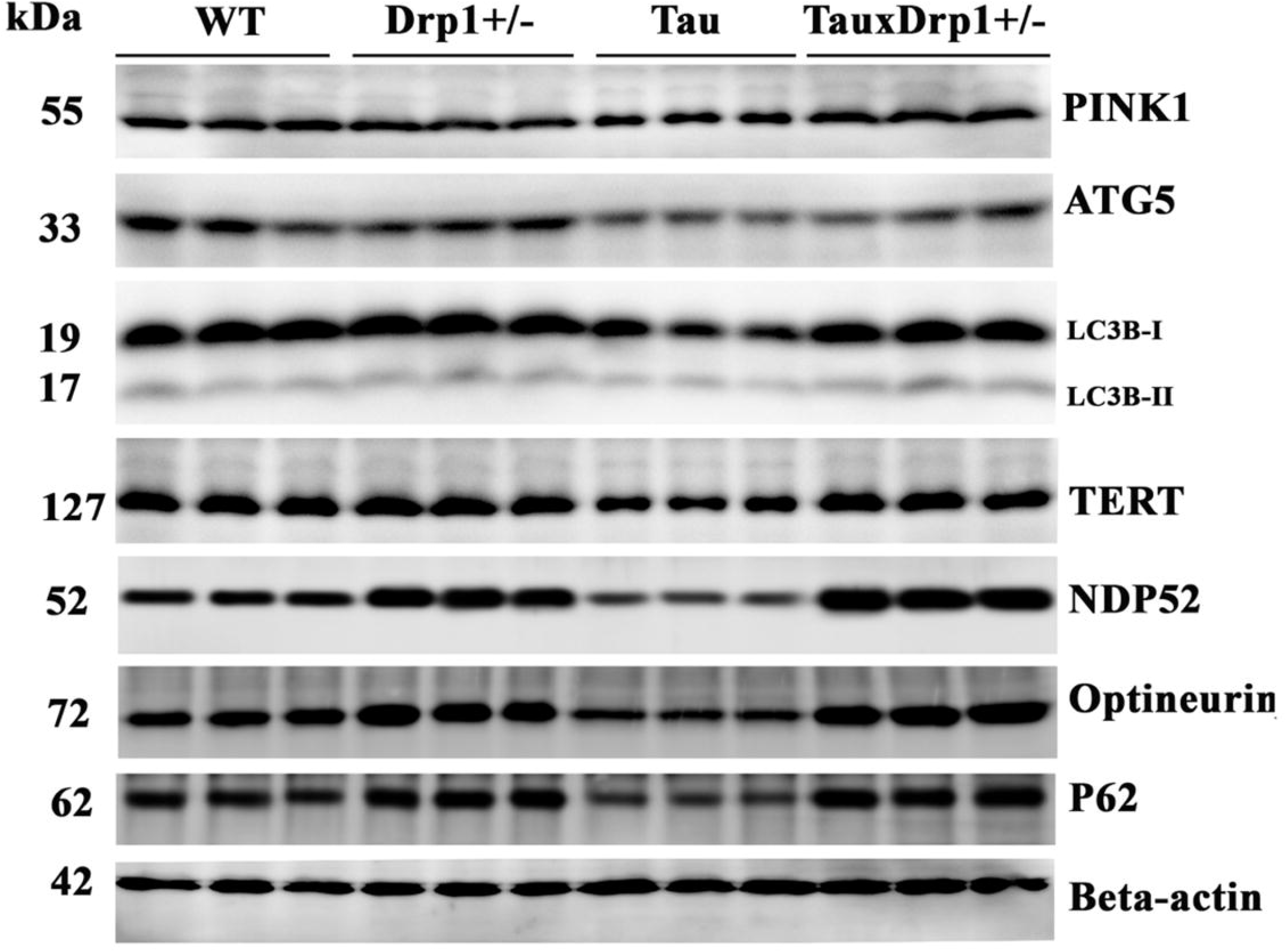
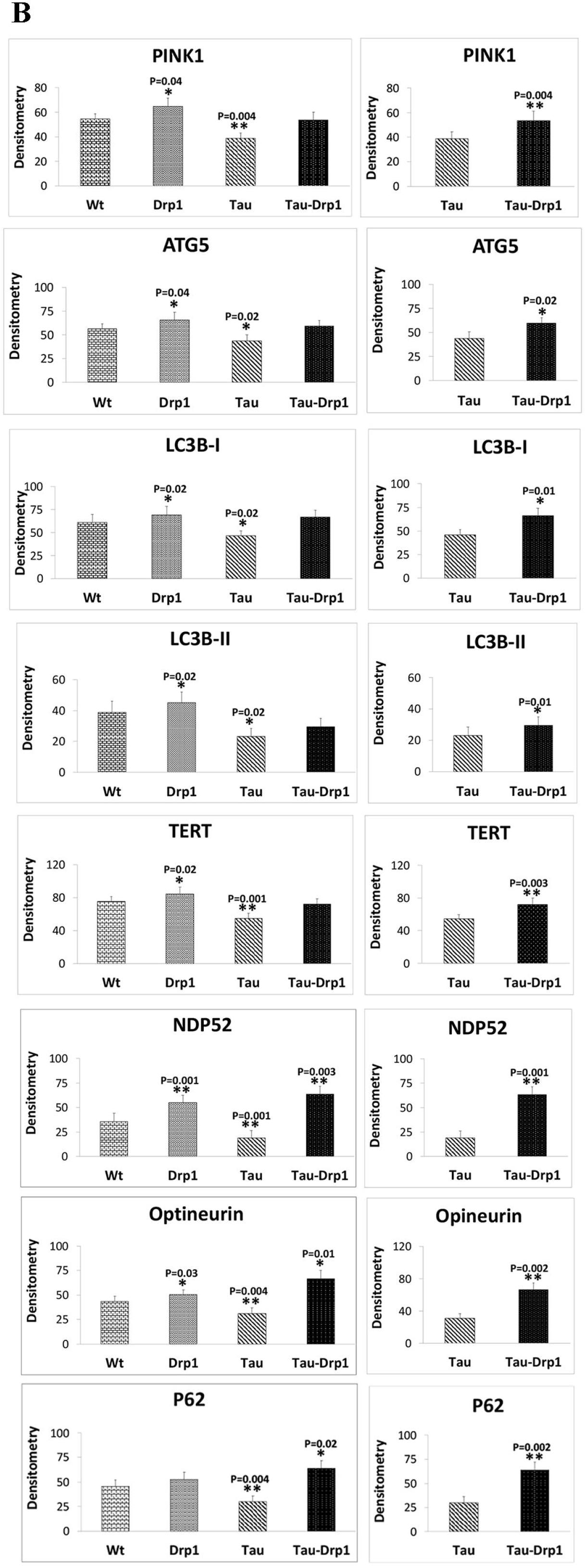
Immunoblotting analysis of autophagy and mitophagy proteins. Immunoblotting analysis was performed for autophagy proteins ATG5, LC3BI, LC3BII, P62, TERT (autophagy inducer) in 12-month-old WT, Drp1+/-m Tau, and double mutant Drp1+/- X Tau mice. Autophagy proteins ATG5, (P=0.02) LC3BI (P=0.02), LC3BII (P=0.02) and P62, LC3-binding adaptor protein (P=0.004), and TERT (autophagy inducer) (P=0.001) were significantly decreased in 12-month-old Tau mice relative to WT mice. Mitophagy proteins, PINK1 (P=0.004), Optineurin (P=0.004), and NDP52 (0.001) (mitophagy receptor proteins) were also significantly reduced in Tau mice relative control WT mice. Mitophagy proteins (P=0.04), optineurin (P=0.03), and NDP52 (0.001) (mitophagy receptor proteins) were also significantly increased in Drp1+/- mice relative control, WT mice. On the other hand, autophagy proteins ATG5, (P=0.04) LC3B-I (P=0.02), LC3B-II (P=0.02), and TERT (autophagy inducer) (P=0.02) were significantly decreased in 12-month-old Tau mice relative to age-matched WT mice. Protein levels P62 were increased, but not significant in Drp1+/- mice relative to WT mice. In comparison to Tau mice, Drp1+/- X Tau mice autophagy proteins ATG5, (P=0.02) LC3B-I (P=0.01), LC3-BII (P=0.01) and P62 (P=0.002) and TERT (autophagy inducer) (P=0.001) were significantly increased. Mitophagy proteins, PINK1 (P=0.004), Optineurin (P=0.002), and NDP52 (0.001) (mitophagy receptor proteins) were also significantly increased in Drp1+/- X Tau mice relative to age-matched Tau mice.

##### Drp1+/- and WT mice

As shown in **Figure 2**, mitophagy proteins (P=0.04), optineurin (P=0.03), and NDP52 (0.001) (mitophagy receptor proteins) were also significantly increased in Drp1+/- mice relative control, WT mice. On the other hand, autophagy proteins ATG5, (P=0.04) LC3BI (P=0.02), LC3BII (P=0.02), and TERT (autophagy inducer) (P=0.02) were significantly decreased in 12-month-old Tau mice relative to age-matched WT mice. Protein levels P62 were increased, but not significant in Drp1+/- mice relative to WT mice.

##### Drp1+/- X Tau and Tau mice

As shown in **Figure 2**, autophagy proteins ATG5, (P=0.02) LC3B-I (P=0.01), LC3B-II (P=0.01) and P62 (P=0.002) and TERT (autophagy inducer) (P=0.001) were significantly increased in 12-month-old Drp1+/- X Tau mice relative to age-matched Tau mice. Mitophagy proteins, PINK1 (P=0.004), Optineurin (P=0.002), and NDP52 (0.001) (mitophagy receptor proteins) were also significantly increased in Drp1+/- X Tau mice relative to age-matched Tau mice.

#### Mitochondrial Biogenesis proteins

To determine the protective effects of a partial reduction of Drp1, we assessed mitochondrial biogenesis proteins, PGC1α, Nrf1, Nrf2, and TFAM in 12-month-old WT, Drp1, Tau, and Drp1+/- X Tau mice.

##### Tau and WT mice

As shown in **Figure 3**, mitochondrial biogenesis proteins PGC1a, (P=0.04) Nrf1 (P=0.01), and TFAM (P=0.01) were significantly decreased in 12-month-old Tau mice relative to age-matched WT mice.

**Figure 3.**
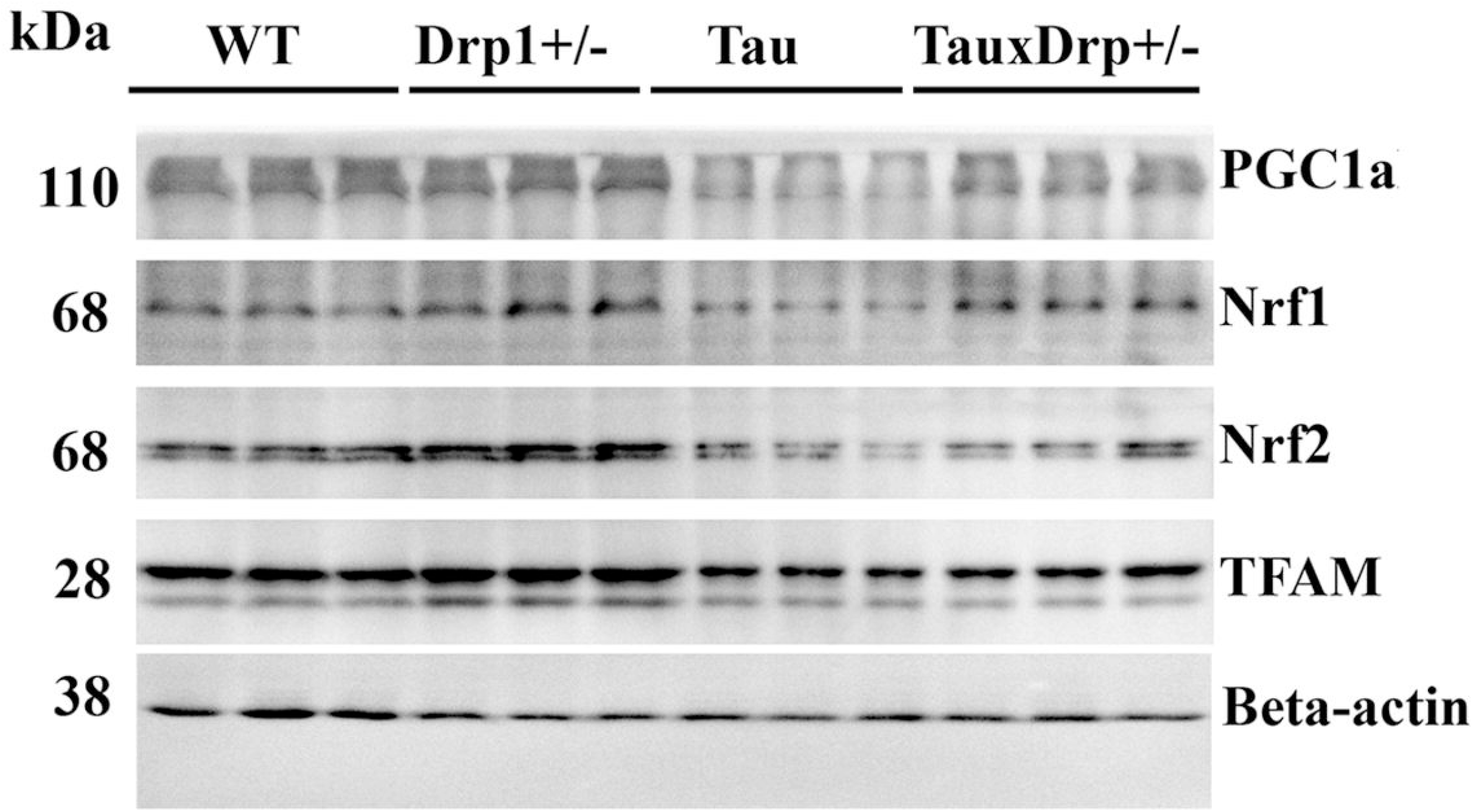
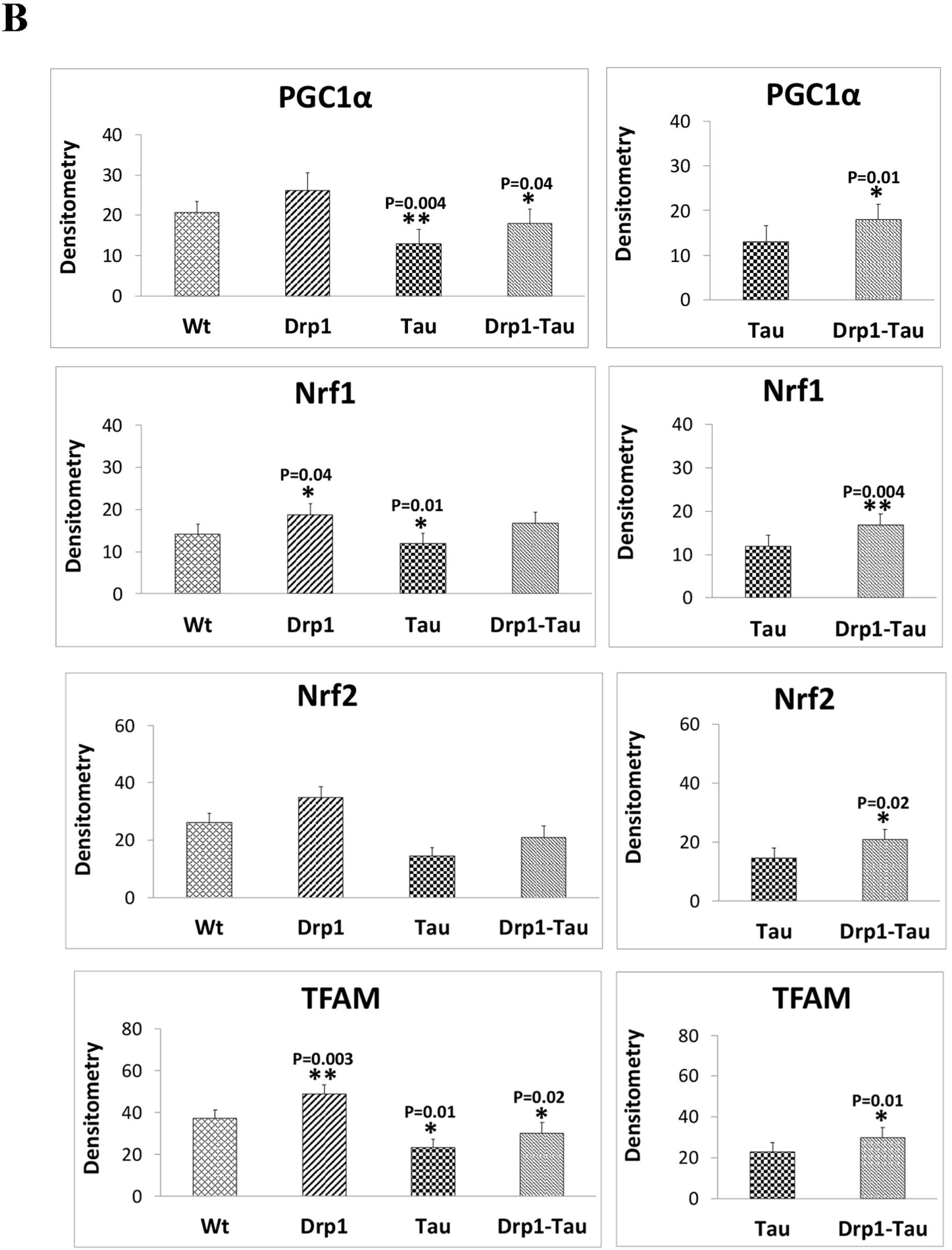
Immunoblotting analysis of mitochondrial Biogenesis proteins in 12-month-old WT, Drp1, Tau, and Drp1+/- X Tau mice. Significantly decreased mitochondrial biogenesis proteins PGC1a, (P=0.04) Nrf1 (P=0.01), and TFAM (P=0.01) were found in 12-month-old Tau mice relative to age-matched WT mice. These were significantly increased in Drp1+/- mice relative to WT mice. PGC1a and Nrf2 were increased Drp1+/- mice, but not significant. In double mutant mice relative to Tau mice, mitochondrial biogenesis proteins PGC1a, (P=0.041) Nrf1 (P=0.004), Nrf2 (P=0.02), and TFAM (P=0.01) were significantly increased.

##### Drp1+/- and WT mice

Mitochondrial biogenesis proteins, Nrf1 (P=0.04), and TFAM (P=0.003) were significantly increased in Drp1+/- mice relative control, WT mice. PGC1α and Nrf2 were increased Drp1+/- mice, but not significant (**Figure 3)**.

##### Drp1+/- X Tau and Tau mice

As shown in Figure, all mitochondrial biogenesis proteins PGC1a, (P=0.041) Nrf1 (P=0.004), Nrf2 (P=0.02) and TFAM (P=0.01) were significantly increased in 12-month-old Drp+/- X Tau mice relative to age-matched Tau mice (**Figure 3**).

#### Mitochondrial Dynamics Proteins

We also assessed mitochondrial dynamics proteins in 12-month-old WT, Drp1, Tau, and Drp1+/- X Tau mice.

##### Tau and WT mice

As shown in **Figure 4**, mitochondrial fission proteins Drp1 (P=0.003) and Fis1 (0.01) were significantly increased; and the fusion proteins Mfn1 (P=0.002), Mfn2 (P=0.01), and Opa1 (P=0.01) were significantly decreased in 12 months old Tau mice relative to WT mice, indicating mitochondrial fragmentation.

**Figure 4.**
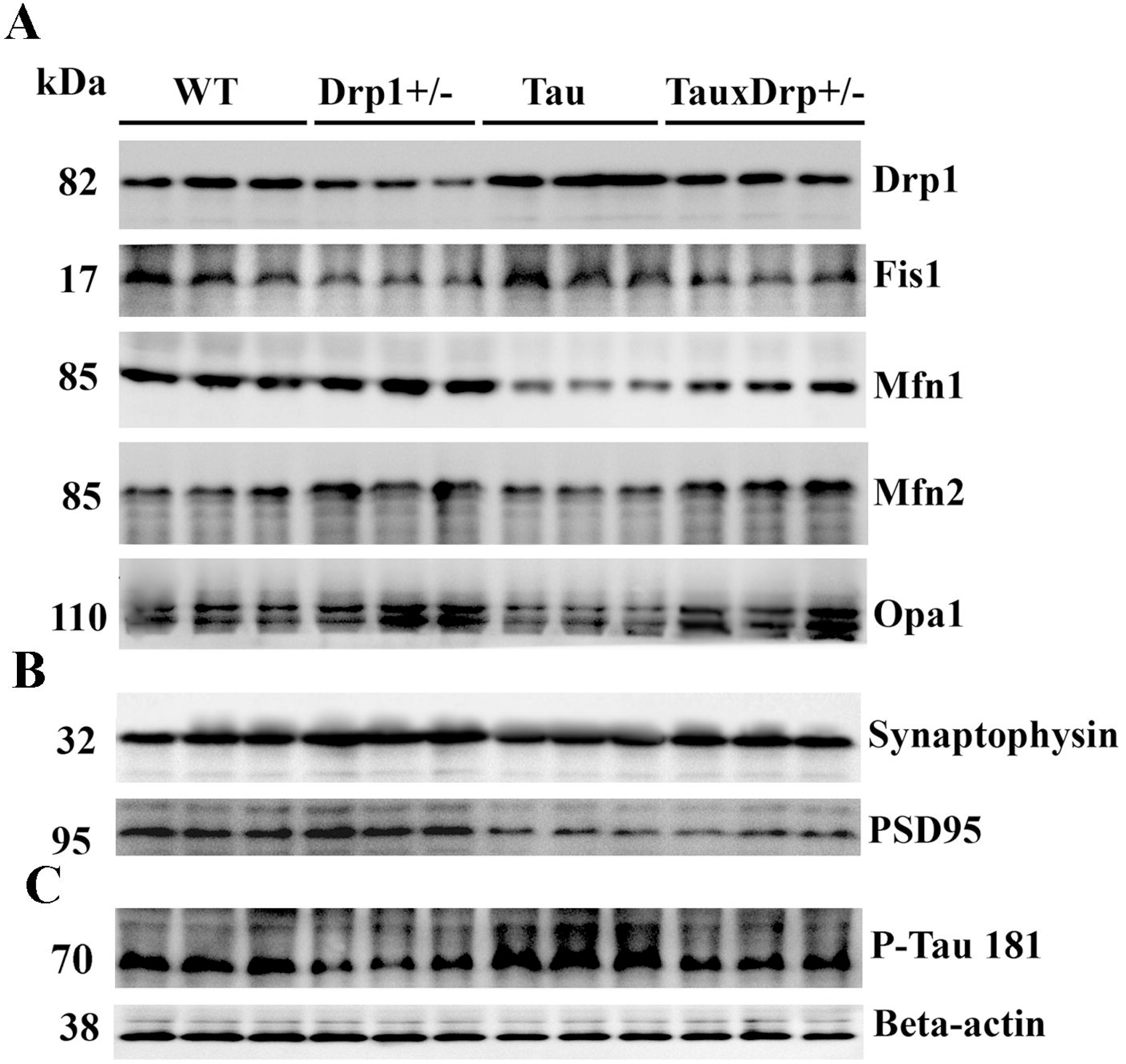
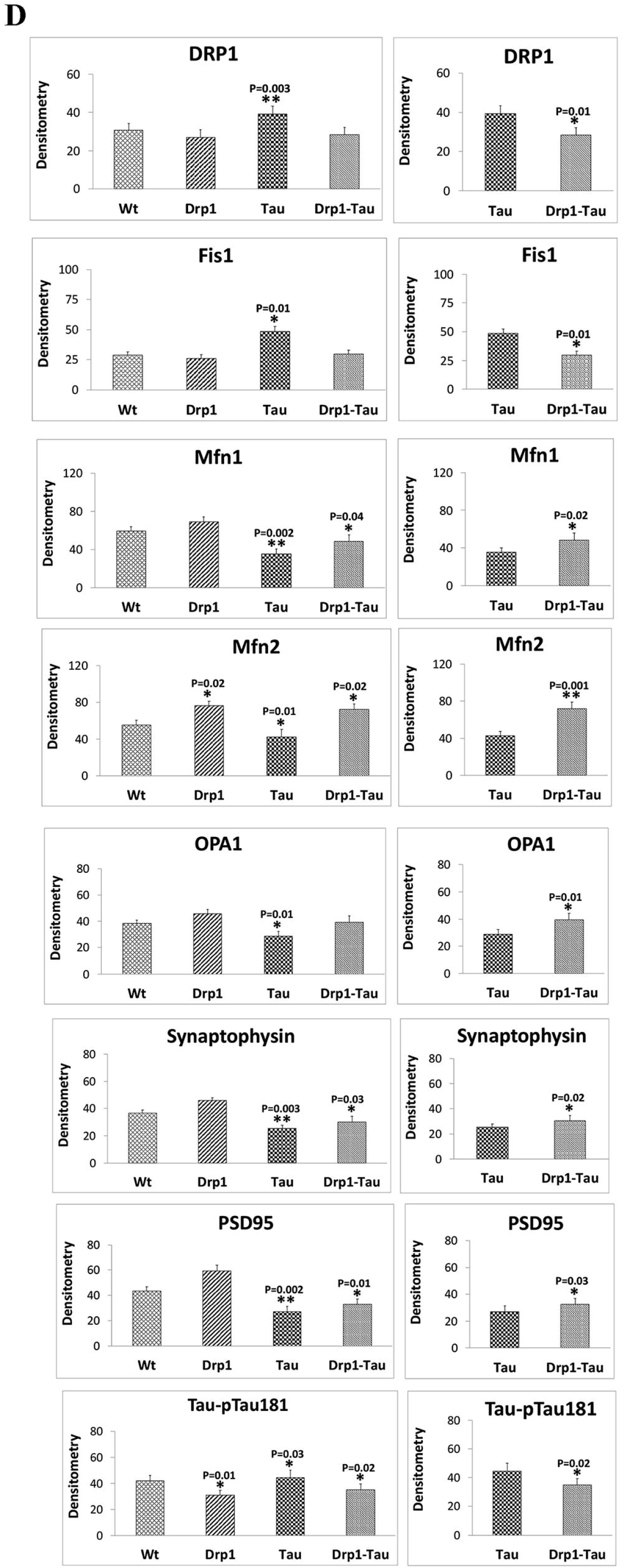
Immunoblotting analysis of mitochondrial dynamics proteins in 12-month-old WT, Drp1, Tau and Drp1+/- X Tau mice. **(A)**. Shows mitochondrial dynamics proteins, Drp1 (P=0.003) and Fis1 (0.01) were significantly increased; and the fusion proteins Mfn1 (P=0.002), Mfn2 (P=0.01), and Opa1 (P=0.01) were significantly decreased in 12 months old Tau mice relative to WT mice, indicating mitochondrial fragmentation in Tau mice. On the contrary, the fission proteins Drp1 (P=0.01) and Fis1 (P=0.01) were significantly decreased and the fusion proteins Mfn1 (P=0.01), Mfn2 (0.04), and Opa1 (P=0.01) were significantly increased 12-month-old double mutant (Drp1+/- X mice relative to Tau mice (P, 0.0001). **(B)** Levels of synaptic proteins synaptophysin (P=0.003), and PSD95 (P=0.002) were significantly reduced in 12-month-old Tau mice. Significantly increased synaptic proteins synaptophysin (P=0.02) and PSD95 (P=0.03) were found in 12-month-old double mutant (Drp1+/- X Tau mice) relative to Tau mice. **(C)** Hyperphosphorylated Tau181 protein levels were increased in Tau mice (P=0.03) relative to WT mice. On the other hand, P-Tau181 levels were reduced Drp1+/- mice (P=0.01) (Figure 4). As expected, P-Tau181 levels were significantly reduced in double mutant (Drp1+/- X Tau) mice relative to Tau mice.

##### Drp1+/- X Tau and Tau mice

On the contrary, the fission proteins Drp1 (P=0.01) and Fis1 (P=0.01) were significantly decreased (**Figure 4**); and the fusion proteins Mfn1 (P=0.01), Mfn2 (0.04), and Opa1 (P=0.01) were significantly increased 12-month-old Tau-Drp1 mice relative to Tau mice (P, 0.0001), indicating that reduced Drp1 protects against phosphorylated Tau-induced mitochondrial dynamics toxicity.

#### Synaptic Proteins

We measured synaptic proteins, synaptophysin, and PSD95 in 12-month-old WT, Drp1, Tau, and Drp1+/- X Tau mice, to understand the impact of reduced Drp1 in Tau mice **(Figure 4)**.

##### Tau and WT mice

As shown in Figure, the levels of synaptic proteins synaptophysin (P=0.003), and PSD95 (P=0.002) were significantly reduced in 12-month-old Tau mice (**Figure 4**).

##### Drp1+/- X Tau and Tau

Synaptic proteins synaptophysin (P=0.02) and PSD95 (P=0.03) were significantly increased in the 12-month-old double mutant (Drp1+/- X Tau mice) relative to Tau mice (**Figure 4**).

#### Hyperphosphorylated Tau181 protein

We also assessed hyperphosphorylated tau 181 protein in all 4 lines of mice. As shown in Figure P-Tau181 levels were increased in Tau mice (P=0.03) relative to WT mice. On the other hand, P-Tau181 levels were reduced in Drp1+/- mice (P=0.01) (**Figure 4**). As expected, P-Tau181 levels were significantly reduced in double mutant (Drp1+/- X Tau) mice relative to Tau mice.

### Mitochondrial functional parameters

Mitochondrial function was assessed in 12-month-old WT, Drp1+/-, Tau, and Tau-Drp1 mice, to understand the impact of reduced Drp1 on mitochondrial ATP, H_2_O_2_ production, lipid peroxidation (4-hydroxy-nonenol - HNE), and cytochrome c oxidase activity.

#### WT and Tau

Compared to 12-month-old WT mice, in Tau mice, the levels of hydrogen peroxide (P=0.005) and 4-hydroxy-nonenol (P=0.002) were significantly higher, while the levels of cytochrome oxidase (P=0.01) and ATP (P=0.001) (D) were significantly lower (**Figure 5**).

**Figure 5.**
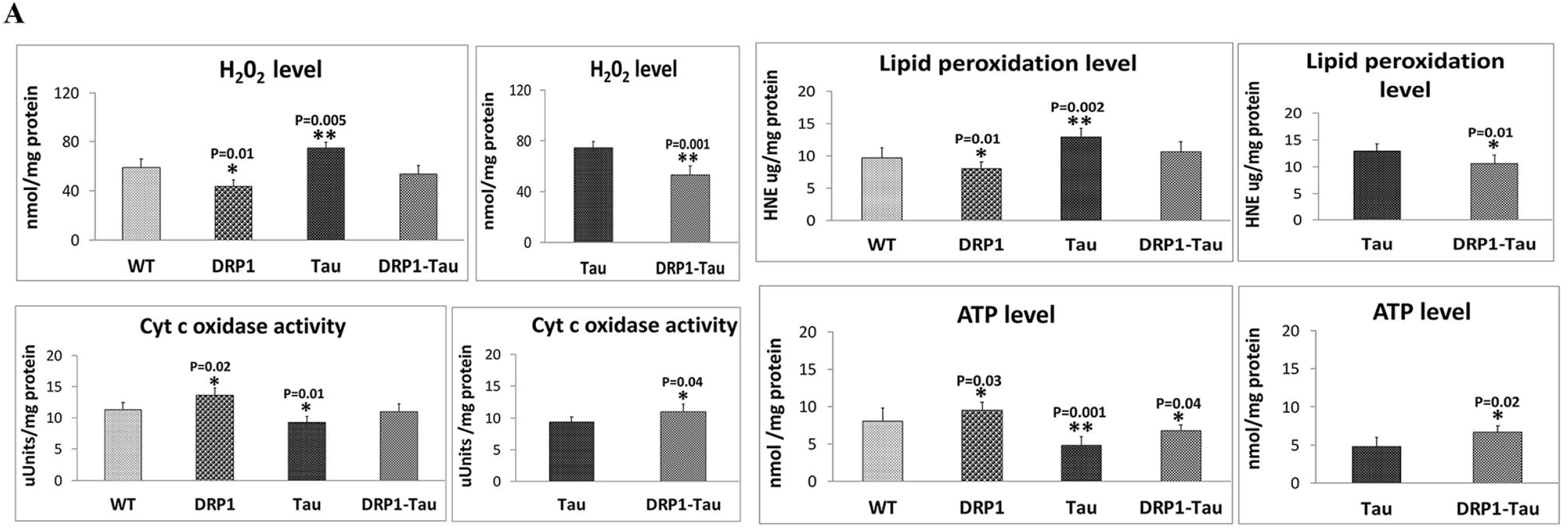
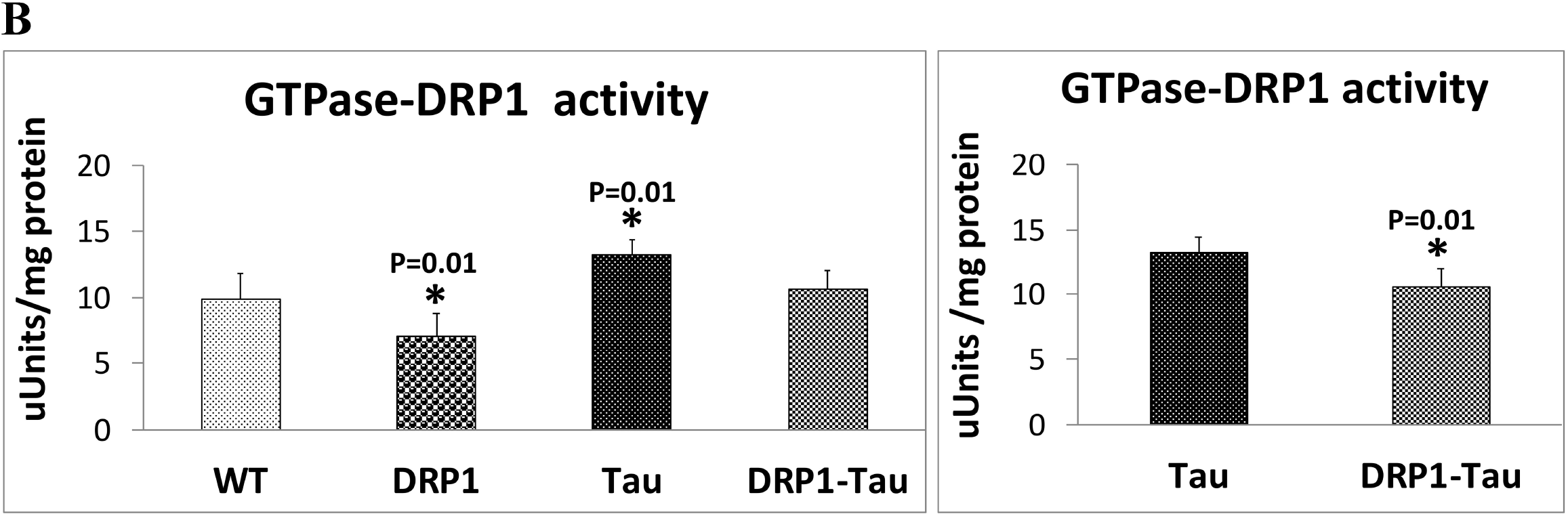
Mitochondrial functional parameters in 12-month-old TW, Drp1, Tau, and Tau-Drp1 mice. **(A) Hydrogen peroxide production, lipid peroxidation (4-HNE), cytochrome oxidase activity and ATP levels.** Compared to 12-month-old WT mice, in Tau mice, the levels of hydrogen peroxide (P=0.005) and 4-hydroxy-nonenol (P=0.002) were significantly higher, while the levels of cytochrome oxidase (P=0.01) and ATP (P=0.001) were significantly lower. The levels of hydrogen peroxide (P=0.01) and 4-hydroxy-nonenol (P=0.02) levels were significantly decreased and cytochrome c oxidase activity (P=0.02) and mitochondrial ATP (P=0.003) were significantly increased in 12-month-old Drp1+/- mice relative to age-matched WT mice. Both hydrogen peroxide (P=0.001) and 4-hydroxy-nonenol (P=0.01) levels significantly increased in 12-month-old Drp1 X Tau mice relative to age-matched Tau mice. Overall, mitochondrial function is defective in Tau mice and defective mitochondrial function is reduced in a partial reduction of Drp1 in Tau mice. **(B) GTPase Drp1 enzymatic activity** was assessed GTPase Drp1. Significantly increased GTPase Drp1 activity (P=0.01) was observed in 12-month-old Tau mice relative to age-matched WT mice. GTPase Drp1 activity was significantly reduced in Drp1+/- mice (P=0.01) relative to WT mice. In comparison to Tau mice, GTPase Drp1 activity was significantly reduced in Drp1+/- X Tau mice (P=0.01). These observations strongly suggest that reduced GTPase Drp1 activity was correlated with decreased mitochondrial fragmentation in double mutant mice

#### WT and Drp1+/-

As shown in **Figure 5**, hydrogen peroxide (P=0.01) and 4-hydroxy-nonenol (P=0.02) levels were significantly decreased and cytochrome c oxidase activity (P=0.02) and ATP (P=0.003) were significantly increased in 12-month-old Drp1+/- mice relative to age-matched WT mice.

#### Drp1 and Drp1 X Tau

Both hydrogen peroxide (P=0.001) and 4-hydroxy-nonenol (P=0.01) levels significantly increased in 12-month-old Drp1 X Tau mice relative to age-matched Tau mice (**Figure 5**).

Overall, these observations strongly suggest that mitochondrial function is defective in Tau mice and defective mitochondrial function is reduced in a partial reduction of Drp1 in Tau mice.

### GTPase Drp1 enzymatic activity

We also assessed GTPase Drp1 enzymatic activity, to understand the correlation between increased mitochondrial fragmentation GTPase Drp1 enzymatic activity in Tau mice and the impact of reduced Drp1 in mitochondrial fragmentation. As shown in **Figure 5**, significantly increased GTPase Drp1 activity (P=0.01) was observed in 12-month-old Tau mice relative to age-matched WT mice.

As expected, significantly decreased levels of GTPase Drp1 activity were observed in Drp1+/- mice (P=0.01) relative to WT mice. In comparison to Tau mice, GTPase Drp1 activity was significantly reduced in Drp1+/- X Tau mice (P=0.01) (**Figure 5**).

These observations strongly suggest that reduced GTPase Drp1 activity was correlated with mitochondrial fragmentation, in other words, increased GTPase Drp1 activity enhances mitochondrial fragmentation and reduced Drp1 activity decreases mitochondrial fragmentation.

### Transmission Electron Microscopy

To determine the impact of reduced Drp1 on mitochondrial number and length in 12-month-old non-transgenic WT mice, Tau (P301L) mice, Drp1+/- mice, and double mutant (Drp1+/- X Tau) mice, we performed transmission electron microscopy in hippocampal and cortical tissues from 12-month-old non-transgenic WT mice, Tau, Drp1+/-, and double mutant (Drp1 X Tau) mice.

#### Mitochondrial number-hippocampus

In comparison to age-matched WT mice, Drp1+/- mice (P=0.01), showed a reduced number of mitochondria (**Figure 6A**). On the other hand, Tau mice showed a significantly increased number of mitochondria (P=0.002) relative to WT mice, indicating that mitochondrial fragmentation is increased in Tau mice. In comparison to Tau mice, double mutant (Drp1 X Tau) mice showed a significantly reduced number of mitochondria, indicating reduced Drp1 decreased mitochondrial number.

**Figure 6:**
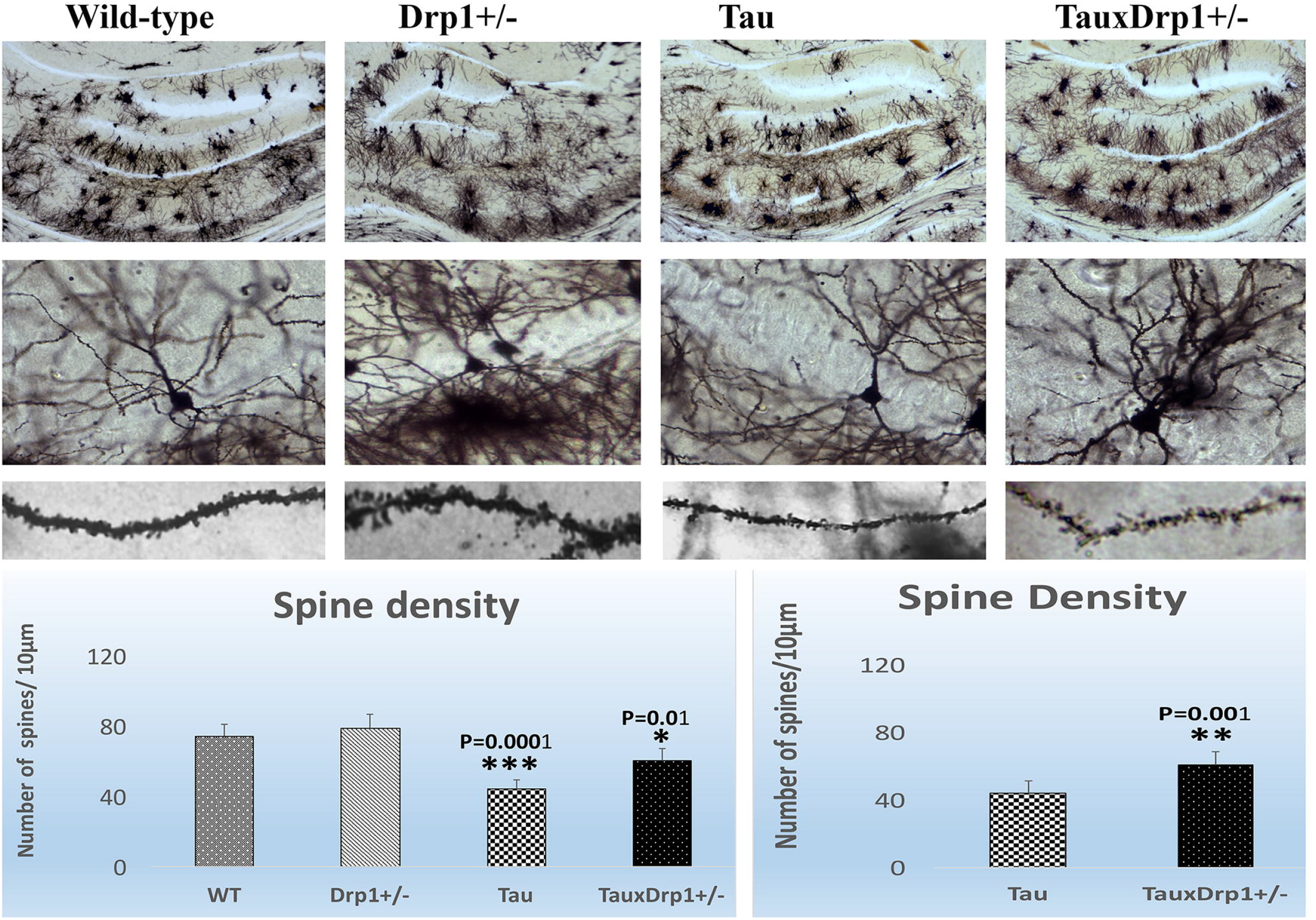
Transmission electron microscopy analysis. Using transmission electron microscopy, we assessed mitochondrial number and length in hippocampal and cortical tissues from 12-month-old non-transgenic WT mice, Tau, Drp1+/-, and double mutant (Drp1 X Tau) mice. In comparison to age-matched WT mice, Tau mice (P=0.002), showed an increased number of mitochondria in the hippocampus (**Figure 6A**). On the other hand, Drp1+/- mice showed a significantly reduced number of hippocampal mitochondria (P=0.01) relative to WT mice. In comparison Tau mice, double mutant (Drp1 X Tau) mice showed a significantly reduced number of hippocampal mitochondria, indicating reduced Drp1 decreases fission activity. The significantly reduced cortical mitochondrial number is observed in 12-month-old Drp1+/- mice to age-matched WT mice, Drp1+/- mice (P=0.01) (**Figure 6B**). In comparison to 12-month-old WT mice, Tau mice showed a significantly increased number of cortical mitochondria (P=0.002). A significantly reduced number of cortical mitochondria is found in double mutant (Drp1 X Tau) mice relative to Tau mice. The hippocampal mitochondrial length was significantly decreased in 12-month-old Tau mice relative to WT mice (P=0.004) (**Figure 6A**). On the other hand, significantly increased hippocampal mitochondrial length was found in 12-month-old Drp1 X Tau mice relative to Tau mice (P=0.001). The significantly reduced cortical mitochondrial length was observed in 12-month-old Drp1+/- mice to age-matched WT mice, Drp1+/- mice (P=0.005) (**Figure 6B**). A significantly increased cortical mitochondrial length was found in Drp1 X Tau mice relative to Tau mice (P=0.003).

#### Mitochondrial number-cortex

Significantly reduced mitochondrial number is observed in 12-month-old Drp1+/- mice to age-matched WT mice, Drp1+/- mice (P=0.01) (**Figure 6B**). In comparison to 12-month-old WT mice, Tau mice showed a significantly increased number of mitochondria (P=0.002). A significantly reduced number of mitochondria is found in double mutant (Drp1 X Tau) mice relative to Tau.

These observations suggest that increased mitochondrial fragmentation in both hippocampi and cortices of Tau mice and these were reduced in double mutant (Drp1 X Tau) mice, in other words, a partial reduction of Drp1 reduces excessive fragmentation of mitochondria.

#### Mitochondrial length-hippocampus

Mitochondrial length was significantly decreased in 12-month-old Tau mice relative to WT mice (P=0.004) (**Figure 6A**). On the other hand, significantly increased mitochondrial length was found in 12-month-old Drp1 X Tau mice relative to Tau mice (P=0.001).

#### Mitochondrial length-cortex

Significantly reduced mitochondrial length was observed in 12-month-old Drp1+/- mice to age-matched WT mice, Drp1+/- mice (P=0.005) (**Figure 6B**). A significantly increased mitochondrial length was found in Drp1 X Tau mice relative to Tau mice (P=0.003).

These observations indicate that phosphorylated tau enhances fragmentation of mitochondria, on the contrary, a partial reduction of Drp1 increases mitochondrial length in Tau mice.

### Golgi-Cox Staining and Dendritic spine density

To determine the impact of reduced Drp1+/- on dendritic spines, we assessed dendritic spines in the hippocampal neurons of 12-month-old all 4 lines of mice - WT mice, Tau mice, Drp1+/- mice, and double mutant (Drp1+/- X Tau) mice. As shown in **Figure 7**, in comparison to WT mice, Tau mice had a significantly reduced number of dendritic spines (P=0.0001). In double mutant (Drp1 X Tau) mice, spine density was significantly higher (P=0.001) than in Tau mice (**Figure 7**).

**Figure 7.**
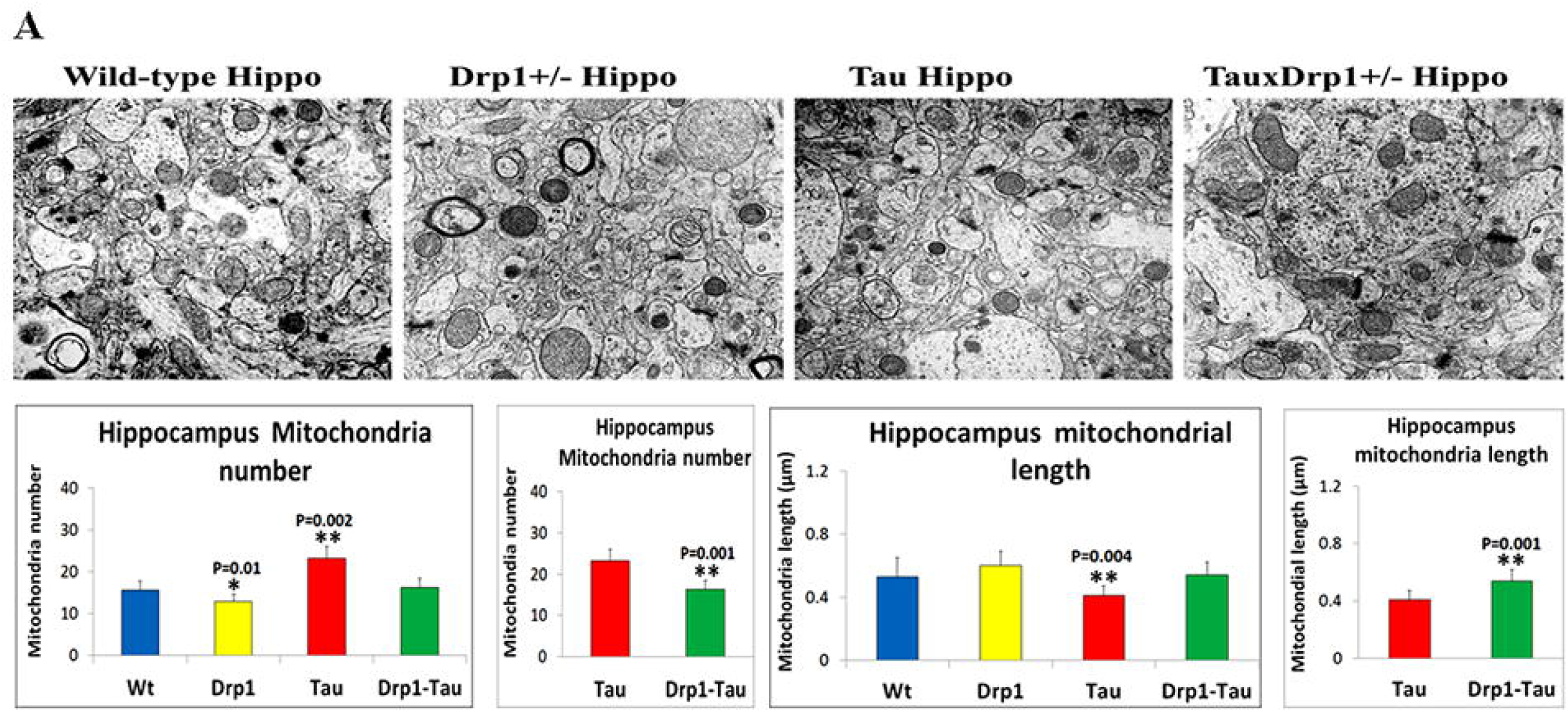
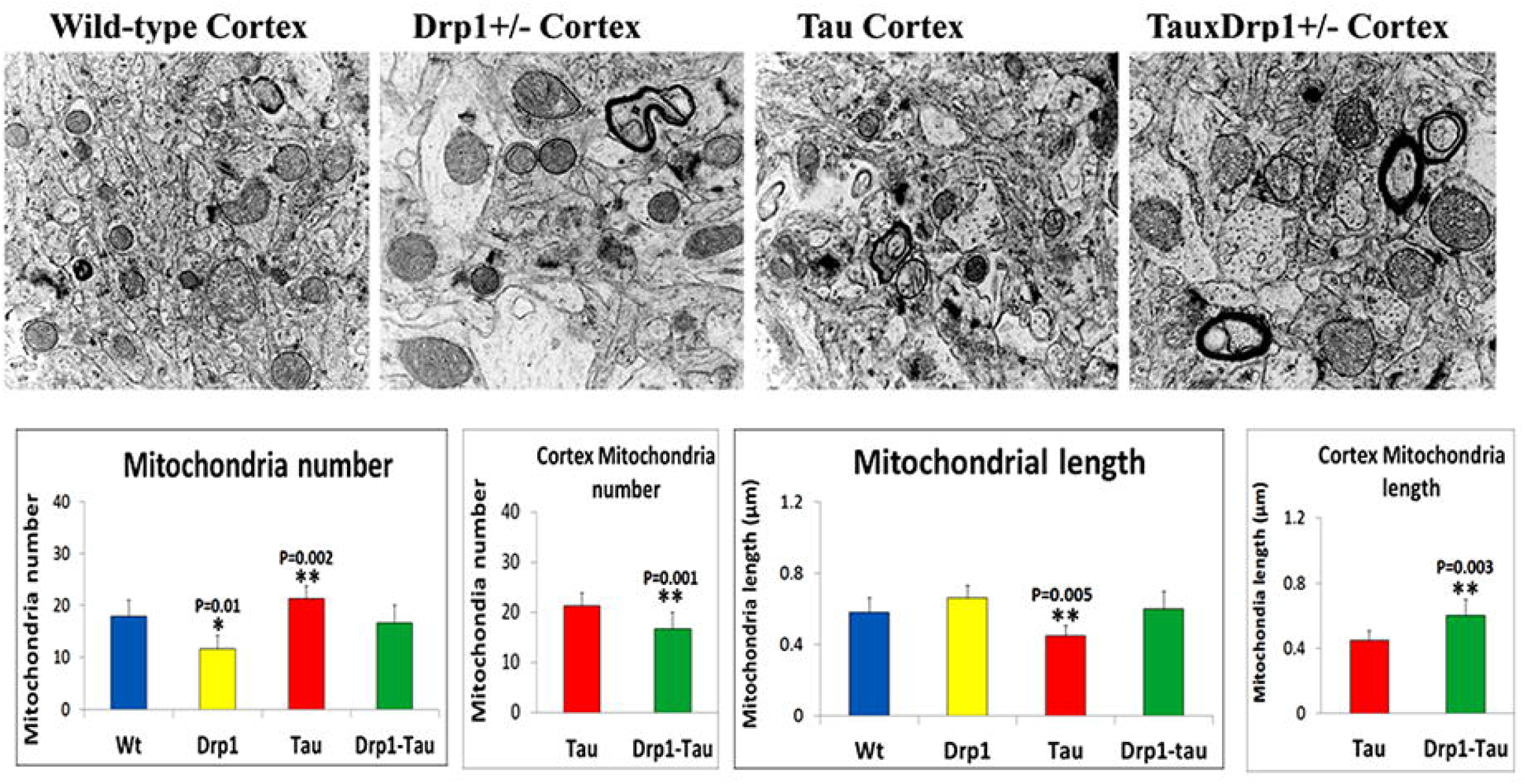
Dendritic spine analysis using Golgi-cox staining in 12-month-old non-transgenic wild-type mice, Tau mice, Drp1+/- mice, and double mutant (Drp1+/- X Tau) mice. **(A)** Golgi-Cox staining under low magnification (4X). **(B)** Quantification of spine density in WT, Drp1 (+/-), Tau, and double mutant (Drp1-Tau) mice, and (C) enlargement of neuron with dendrites (60X). In comparison to WT mice, Tau mice had significantly fewer dendritic spines (P, 0.0001). In double mutant (Drp1-Tau) mice, spine density was higher (P< 0.001) than in Tau mice, with a p-value of 0.01 when compared to WT mice.

## Discussion

The long-term goal of our study was to understand molecular mechanisms <u>involved in defective</u> mitophagy in the progression and pathogenesis of AD. Emerging evidence from AD cell cultures, mouse and postmortem studies suggest that mitophagy is defective early on in the progression of disease [6, 39, 40, 7, 8, 26, 41, 18, 22, 29, 20, 24, 23]. Several studies provided compelling evidence that Aβ and P-Tau interacts with a large number of mitochondrial proteins such as Drp1, VDAC1 (voltage-dependent anion channel 1), CypD (cyclophilin D) and ABAD (amyloid beta-induced alcohol dehydrogenase), leading to induced free radical production, increased lipid peroxidation, and depletion of major mitophagy proteins, PINK1 and Parkin [6, 39, 40, 7, 8, 26, 41, 42, 22, 29, 20, 24, 23, 43, 44, 45]. These changes ultimately leading to defective clearance of dead or dying mitochondria, synaptic damage.

Our laboratory has been working to understand molecular events in mitochondria, aging, synapse and Aβ & P-Tau in AD over the last 2 decades; we have found that Aβ and P-Tau interacts with mitochondrial fission protein, Drp1, and mitochondrial outer membrane protein VDAC1, these interactions enhance GTPase Drp1 activity, increases mitochondrial fragmentation and most importantly depletes mitophagy proteins PINK1 and Parkin [26, 26, 41, 42, 22, 29, 20, 24, 23]. Based on these observations, we propose that a partial reduction of Drp1 reduces mitochondrial fragmentation, enhances biogenesis, mitophagy/autophagy, synaptic activity, dendritic spines, and improve cognitive functions in AD. To test our hypothesis, we recently crossed Drp1 heterozygote knockout (Drp1+/-) mice with transgenic Tau and transgenic APP mice and studied cellular changes in 6-month-old mice (Manczak et al 2016, Kandimalla et al 2016). Findings from these studies revealed that double mutant (Drp1+/- X Tau; Drp1+/- X APP) mice did better than transgenic APP and transgenic Tau mice, meaning double mutant (Drp1+/- X Tau; Drp1+/- X APP) mice showed reduced mitochondrial fragmentation and increased synaptic and mitochondrial biogenesis proteins. However, in our 6-month-old mice, we did not study cognitive behavior, and assessed mitophagy/autophagy and dendritic spines. In the current study, we assessed cognitive behavior, and assessed mitophagy/autophagy, mitochondrial biogenesis, dynamics and dendritic spines in 12-month-old Drp1+/- X Tau mice relative to Tau mice.

To determine the impact of reduced Drp1 on cognitive behavior in transgenic Tau mice, we performed multiple behavioral tests, including Rotarod and Morris Water Maze tests. When compared to WT, Drp1+/- mice, and double mutant (Drp1+/- X Tau), Tau mice had a considerably longer latency time to detect a platform (**Figure 1**). Drp1+/- mice and double mutant mice resulted in a considerable reduction in the time to identify a hidden platform. In comparison to WT, Drp1+/-, and double mutant (Drp1 X Tau) mice, Tau mice had a slower swimming speed. However, reduced Drp1 enhanced swimming speed considerably. Tau mice spent reduced time (represented in percentage) in all 16 trials on the target quadrant compared with WT, Drp1+/-, and double mutant (Drp1+/- X Tau) mice (**Figure 1**). Overall, Drp1+/- and double mutant mice did better than WT and Tau mice in all behavioral tests studied. Based on these observations, we conclude that a partial reduction of Drp1 improved cognitive behavior in the presence of the Tau mutation.

### mRNA and protein changes

Our extensive analysis of mRNA and protein levels of autophagy, mitophagy, mitochondrial biogenesis, mitochondrial dynamics and synaptic genes in 12-month-old Drp1+/−, Tau, Tau X Drp1+/− mice relative to control WT mice revealed that 1) reduced levels of mRNA and protein levels of all autophagy, mitophagy, biogenesis, synaptic and mitochondrial fusion genes in transgenic Tau mice, 2) however, these were increased in a partial reduction of Drp1 (Drp1+/-) in mice, and 3) our comparative analysis of Drp1+/- X Tau mice and Tau mice showed increased levels of mRNA and protein levels of autophagy, mitophagy, biogenesis, synaptic and mitochondrial fusion genes in a partial reduction of Drp1 in 12-month-old Tau mice. These observations indicate that reduced Drp1 is beneficial to neurons not only in WT but also in Tau mice. Current findings of increased mRNA and protein levels of synaptic, mitochondrial biogenesis, and mitochondrial fusion genes agree with our earlier observations of 6-month-old Drp1+/−, Tau, Drp1+/− X Tau mice [38].

Regarding mitophagy, in the current study, a partial reduction of Drp1 in Tau mice activates PINK1 signaling, with increased levels of PINK1 receptors NDP52 and optineurin with p62. These observations support that Drp1 reduction activates mitophagy and reduces excessive mitochondrial fragmentation in Tau mice. For the first time, we observed a mechanistic link between reduced Drp1 and enhanced autophagy and mitophagy in AD.

In the current study, we also found reduced phosphorylated tau (ser181) protein levels in 12-month-old Drp1+/- and double mutant Drp1 X Tau mice relative to Tau mice and these observations concur with our earlier studies of 6-month-old Drp1+/- and double mutant Drp1 X Tau mice. These findings suggest that reduced Drp1 influences P-Tau levels in the disease state. However, further studies are still needed to understand mechanistic links.

Mitochondrial function (increased ATP production and cytochrome c oxidase activity and decreased free radicals and lipid peroxidation) significantly improved in a partial reduction of Drp1 in both 12-month-old WT and transgenic Tau mice. The levels of GTPase Drp1 enzymatic activity are essential for mitochondrial fission in a healthy state, these were increased AD and enhanced mitochondrial fragmentation [26, 42, 19, 22]. The GTPase Drp1 activity levels were reduced in 12-month-old Drp1+/- and Drp1 X Tau mice and these observations of reduced GTPase Drp1 activity correlate well with mRNA and protein levels of mitochondrial fission genes and reduced excessive mitochondrial fragmentation.

For the first time, using Golgi-cox staining, we assessed dendritic spines in Drp1+/- and Drp1+/- X Tau mice. Our extensive analysis of dendritic spines revealed that increased spine density in 12-month-old Drp1+/- mice relative to WT mice and in 12-month-old double mutant Drp1+/- X Tau mice relative to Tau mice. These observations strongly suggest that reduced Drp1 enhances synaptic activity, mRNA and protein levels of synaptic genes, dendritic spines; and increased synaptic activity and improved cognitive behavior in Drp1+/- mice and Drp1+/- X Tau mice.

Our extensive transmission electron microscopy analysis of hippocampal and cortical tissues revealed that increased mitochondrial length and reduced number of mitochondria in 12-month-old Drp1+/- and Drp1+/- X Tau mice. These observations correlate well with increased mitochondrial function & fusion activity and reduced fission activity in Tau mice. Based on these observations, we cautiously conclude that the quality of mitochondria, mitochondrial health, mitophagy and synaptic activities are improved in a partial reduction of Drp1 in Tau mice.

## References

1. D.J. Selkoe, Alzheimer’s disease: genes, proteins, and therapy. Physiol Rev. 81(2) (2001) 741–66.

2. M.P. Mattson, Pathways towards and away from Alzheimer’s disease. Nature. 430 (7000) (2004) 631–639.

3. F.M. LaFerla, Green KN, Oddo S. Intracellular amyloid-beta in Alzheimer’s disease. Nat Rev Neurosci 8 (7) (2007) 499–509.

4. H. Morton, S. Kshirsagar, E. Orlov, L.E. Bunquin, N. Sawant, L. Boleng, M. George, T. Basu, B. Ramasubramanian, J.A. Pradeepkiran, S. Kumar, M. Vijayan, A.P. Reddy, P.H. Reddy. Defective mitophagy and synaptic degeneration in Alzheimer’s disease: Focus on aging, mitochondria and synapse. Free Radic Biol Med. 172 (2021) 652–667.

5. Association, A. s. Alzheimer’s disease facts and figures. Alzheimer’s & Dementia, 15 (2019) 321–387.

6. K. Hirai, G. Aliev, A. Nunomura, H. Fujioka, R.L. Russell, C.S. Atwood, A.B. Johnson, Y. Kress, H.V. Vinters, M. Tabaton, S. Shimohama, A.D. Cash, S.L. Siedlak, P.L. Harris, P.K. Jones, R.B. Petersen, G. Perry, M.A. Smith. Mitochondrial abnormalities in Alzheimer’s disease. J Neurosci 21 (9) (2001) 3017–3023.

7. X. Wang, B. Su, S.L. Siedlak, P.I. Moreira, H. Fujioka, Y. Wang, G. Casadesus, X. Zhu. Amyloid-beta overproduction causes abnormal mitochondrial dynamics via differential modulation of mitochondrial fission/fusion proteins. Proc Natl Acad Sci U S A. 105 (49) (2008) 19318–19323.

8. X. Wang, B. Su, H.G. Lee, X. Li, G. Perry, M.A. Smith, X. Zhu. Impaired balance of mitochondrial fission and fusion in Alzheimer’s disease. J Neurosci. 29 (28) (2009) 9090–103.

9. R.H. Swerdlow, J.M. Burns, S.M. Khan. The Alzheimer’s disease mitochondrial cascade hypothesis: progress and perspectives. Biochim Biophys Acta. 1842 (8) (2014) 1219–1231.

10. P.H. Reddy, R. Tripathi, Q. Troung, K. Tirumala, T.P. Reddy, V. Anekonda, U.P. Shirendeb, M.J. Calkins, A.P. Reddy, P. Mao, M. Manczak. Abnormal mitochondrial dynamics and synaptic degeneration as early events in Alzheimer’s disease: implications to mitochondria-targeted antioxidant therapeutics. Biochim Biophys Acta. 1822 (5) (2012) 639–649.

11. A. John, P.H. Reddy. Synaptic basis of Alzheimer’s disease: Focus on synaptic amyloid beta, P-tau and mitochondria. Ageing Res Rev. 65 (2021) 101208.

12. J.A. Pradeepkiran, P.H. Reddy. Defective mitophagy in Alzheimer’s disease. Ageing Res Rev. 64 (2020) 101191.

13. C.A. Pope, H.M. Wilkins, R.H. Swerdlow, et al. Mutations in the Amyloid-β Protein Precursor Reduce Mitochondrial Function and Alter Gene Expression Independent of 42-Residue Amyloid-β Peptide. J Alzheimers Dis. 2021 doi: 10.3233/JAD-210366.

14. P.H. Reddy, U.P. Shirendeb. Mutant huntingtin, abnormal mitochondrial dynamics, defective axonal transport of mitochondria, and selective synaptic degeneration in Huntington’s disease. Biochim Biophys Acta.1822 (2012) 101–110.

15. M.J. Calkins, M. Manczak, P. Mao, et al. Impaired mitochondrial biogenesis, defective axonal transport of mitochondria, abnormal mitochondrial dynamics and synaptic degeneration in a mouse model of Alzheimer’s disease. Hum Mol Genet. 20 (2011) 4515–4529.

16. D.A. Butterfield, J. Drake, C. Pocernich, A. Castegna. Evidence of oxidative damage in Alzheimer’s disease brain: central role for amyloid beta-peptide. Trends Mol Med. 7(12) (2001) 548–554.

17. D.A. Butterfield, B. Halliwell. Oxidative stress, dysfunctional glucose metabolism and Alzheimer disease. Nat Rev Neurosci. 20(3) (2019) 148–160.

18. M. Manczak, R. Kandimalla, X. Yin, P.H. Reddy. Hippocampal mutant APP and amyloid beta-induced cognitive decline, dendritic spine loss, defective autophagy, mitophagy and mitochondrial abnormalities in a mouse model of Alzheimer’s disease. Hum Mol Genet. 27(8) (2018) 1332–1342.

19. R. Kandimalla, M. Manczak, X. Yin, R. Wang, P.H. Reddy. Hippocampal phosphorylated tau induced cognitive decline, dendritic spine loss and mitochondrial abnormalities in a mouse model of Alzheimer’s disease. Hum Mol Genet. 27(1) (2018) 30–40.

20. A.P. Reddy, N. Sawant, H. Morton, S. Kshirsagar, L.E. Bunquin, X. Yin, P.H. Reddy. Selective serotonin reuptake inhibitor citalopram ameliorates cognitive decline and protects against amyloid beta-induced mitochondrial dynamics, biogenesis, autophagy, mitophagy and synaptic toxicities in a mouse model of Alzheimer’s disease. Hum Mol Genet. 30(9) (2021) 789–810.

21. M. Manczak, P. Mao, M.J. Calkins, A. Cornea, A.P. Reddy, M.P. Murphy, H.H. Szeto, B. Park, P.H. Reddy. Mitochondria-targeted antioxidants protect against amyloid-beta toxicity in Alzheimer’s disease neurons. J Alzheimers Dis. 20 (Suppl 2): (2010) S609–S6031.

22. P.H. Reddy, M. Manczak, X. Yin, A.P. Reddy. Synergistic Protective Effects of Mitochondrial Division Inhibitor 1 and Mitochondria-Targeted Small Peptide SS31 in Alzheimer’s Disease. J Alzheimers Dis. 62(4) (2018) 1549–1565.

23. S. Kshirsagar, N. Sawant, H. Morton, A.P. Reddy, P.H. Reddy. Protective effects of mitophagy enhancers against amyloid beta-induced mitochondrial and synaptic toxicities in Alzheimer disease. Hum Mol Genet. (2021) ddab262. doi: 10.1093/hmg/ddab262. Epub ahead of print. PMID: 34505123.

24. A.P. Reddy, X. Yin, N. Sawant, P.H. Reddy. Protective effects of antidepressant citalopram against abnormal APP processing and amyloid beta-induced mitochondrial dynamics, biogenesis, mitophagy and synaptic toxicities in Alzheimer’s disease. Hum Mol Genet. 30 (10) (2021) 847–864.

25. X. Wang, M. Allen, S. Li, Z.S. Quicksall, T.A. Patel, T.P. Carnwath, J.S. Reddy, M.M. Carrasquillo, S.J. Lincoln, T.T. Nguyen, K.G. Malphrus, D.W. Dickson, J.E. Crook, Y.W. Asmann, N. Ertekin-Taner. Deciphering cellular transcriptional alterations in Alzheimer’s disease brains. Mol Neurodegener. 15(1) (2020) 38.

26. M. Manczak, M.J. Calkins, P.H. Reddy. Impaired mitochondrial dynamics and abnormal interaction of amyloid beta with mitochondrial protein Drp1 in neurons from patients with Alzheimer’s disease: implications for neuronal damage. Hum Mol Genet. 20 (2011) 2495–509.

27. W. Wang, J. Yin, X. Ma, F. Zhao, S.L. Siedlak, Z. Wang, S. Torres, H. Fujioka, Y. Xu, G. Perry, X. Zhu. Inhibition of mitochondrial fragmentation protects against Alzheimer’s disease in rodent model. Hum Mol Genet. 26 (21) (2017) 4118–4131.

28. S.H. Baek, S.J. Park, J.I. Jeong, S.H. Kim, J. Han, J.W. Kyung, S.H. Baik, Y. Choi, B.Y. Choi, J.S. Park, G. Bahn, J.H. Shin, D.S. Jo, J.Y. Lee, C.G. Jang, T.V. Arumugam, J. Kim, J.W. Han, J.Y. Koh, D.H. Cho, D.G. Jo. Inhibition of Drp1 Ameliorates Synaptic Depression, Aβ Deposition, and Cognitive Impairment in an Alzheimer’s Disease Model. J. Neurosci. 37(20) (2017) 5099–5110.

29. E.F. Fang, Y. Hou, K. Palikaras, et al. Mitophagy inhibits amyloid-β and tau pathology and reverses cognitive deficits in models of Alzheimer’s disease. Nat Neurosci. 22 (2019) 401–412.

30. P.H. Reddy, D.M. Oliver. Amyloid Beta and Phosphorylated Tau-Induced Defective Autophagy and Mitophagy in Alzheimer’s Disease. Cells.8 (2019) 488.

31. J. Wakabayashi, Z. Zhang, N. Wakabayashi, Y. Tamura, M. Fukaya, T.W. Kensler, M. Iijima, H. Sesaki. The dynamin-related GTPase Drp1 is required for embryonic and brain development in mice. J Cell Biol. 186 (6) (2009) 805–816.

32. N. Ishihara, M. Nomura, A. Jofuku, H. Kato, S.O. Suzuki, K. Masuda, H. Otera, Y. Nakanishi, I. Nonaka, Y. Goto, N. Taguchi, H. Morinaga, M. Maeda, R. Takayanagi, S. Yokota, K. Mihara. Mitochondrial fission factor Drp1 is essential for embryonic development and synapse formation in mice. Nat Cell Biol. 11(8) (2009) 958–966.

33. M. Manczak, P.H. Reddy. Abnormal interaction between the mitochondrial fission protein Drp1 and hyperphosphorylated tau in Alzheimer’s disease neurons: implications for mitochondrial dysfunction and neuronal damage. Hum Mol Genet. 21(11): (2012) 2538-2547.

34. M. Manczak, R. Kandimalla, D. Fry, H. Sesaki, P.H. Reddy. Protective effects of reduced dynamin-related protein 1 against amyloid beta-induced mitochondrial dysfunction and synaptic damage in Alzheimer’s disease. Hum Mol Genet. 25 (23) (2016) 5148–5166.

35. R. Kandimalla, P.H. Reddy. Multiple faces of dynamin-related protein 1 and its role in Alzheimer’s disease pathogenesis. Biochim Biophys Acta. 1862 (4) (2016) 814–828.

36. M .Manczak, P.H. Reddy. Abnormal interaction of oligomeric amyloid-β with phosphorylated tau: implications to synaptic dysfunction and neuronal damage. J Alzheimers Dis. 36 (2) (2013) 285–295.

37. J. Lewis, E. McGowan, J. Rockwood, H. Melrose, P. Nacharaju, M. Van Slegtenhorst, K. Gwinn-Hardy, M. Paul Murphy, M. Baker, X. Yu, K. Duff, J. Hardy, A. Corral, W.L. Lin, S.H. Yen, D.W. Dickson, P. Davies, M. Hutton. Neurofibrillary tangles, amyotrophy and progressive motor disturbance in mice expressing mutant (P301L) tau protein. Nat Genet. 25(4) (2000) 402–405.

38. R. Kandimalla, M. Manczak, D. Fry, Y. Suneetha, H. Sesaki, P.H. Reddy. Reduced dynamin-related protein 1 protects against phosphorylated Tau-induced mitochondrial dysfunction and synaptic damage in Alzheimer’s disease. Hum Mol Genet. 25(22) (2016) 4881–4897.

39. P.H. Reddy, S. McWeeney, B.S. Park, M. Manczak, R.V. Gutala, D. Partovi, Y. Jung, V. Yau, R. Searles, M. Mori, J. Quinn. Gene expression profiles of transcripts in amyloid precursor protein transgenic mice: up-regulation of mitochondrial metabolism and apoptotic genes is an early cellular change in Alzheimer’s disease. Hum Mol Genet 13 (12) (2004) 1225–1240.

40. M. Manczak, T.S. Anekonda, E. Henson, B.S. Park, J. Quinn, P.H. Reddy. Mitochondria are a direct site of A beta accumulation in Alzheimer’s disease neurons: implications for free radical generation and oxidative damage in disease progression. Hum Mol Genet. 15 (9) (2006) 1437–1449.

41. X. Ye, X. Sun, V. Starovoytov, Q. Cai. Parkin-mediated mitophagy in mutant hAPP neurons and Alzheimer’s disease patient brains. Hum Mol Genet.24(10) (2015) 2938–2951.

42. M. Manczak, R. Kandimalla, X. Yin, P.H. Reddy. Hippocampal mutant APP and amyloid beta-induced cognitive decline, dendritic spine loss, defective autophagy, mitophagy and mitochondrial abnormalities in a mouse model of Alzheimer’s disease. Hum Mol Genet. 27(8) (2018) 1332–1342.

43. Q. Cai, H.M. Zakaria, A. Simone, Z.H. Sheng. Spatial parkin translocation and degradation of damaged mitochondria via mitophagy in live cortical neurons. Curr Biol. 22(6) (2012) 545–552.

44. Q. Cai, H.M. Zakaria, Z.H. Sheng. Long time-lapse imaging reveals unique features of PARK2/Parkin-mediated mitophagy in mature cortical neurons. Autophagy. 8(6) (2012) 976–978.

45. Q. Cai, Y.Y. Jeong. Mitophagy in Alzheimer’s Disease and Other Age-Related Neurodegenerative Diseases. Cells. 9(1) (2020) 150.

